# Genetic compensation in *podocalyxin-like* mutants during zebrafish liver development

**DOI:** 10.1101/2025.04.07.647300

**Authors:** Alexis N. Ross, Natalie M. Miscik, Sharanya Maanasi Kalasekar, James Harris, Mimi Tran, Aavrati Saxena, Steven Andrew Baker, Kimberley Jane Evason

## Abstract

Hepatic stellate cells (HSCs) are critical for normal liver development and regeneration. *Podocalyxin-like (podxl)* is highly expressed in zebrafish HSCs, but its role in liver development is not known. Here we report that *podxl* knockdown using CRISPR/Cas9 (“CRISPants”) significantly decreased HSC number in zebrafish larvae at different time points and in two independent HSC reporter lines, supporting a role for *podxl* in HSC development. We generated five *podxl* mutants, including two mutants lacking the predicted *podxl* promoter region, and found that none of the mutants recapitulated the knockdown phenotype. *Podxl* CRISPR/Cas9 injection in mutants lacking the *podxl* guide RNA cut site did not affect HSC number, supporting the hypothesis that the CRISPant phenotype was specific, requiring intact *podxl. Podxl* mRNA levels in three *podxl* mutants were similar to those of wildtype controls. RNA sequencing of *podxl* mutants and controls showed no significant change in transcript levels of genes with sequence similarity to *podxl*, but it revealed upregulation of a network of extracellular matrix genes in *podxl* mutants. These results support a role for *podxl* in zebrafish liver development and suggest that upregulation of a group of functionally related genes represents the main mechanism of compensation for *podxl* genomic loss.

## Introduction

Hepatic stellate cells (HSCs) are mesenchymal cells in the liver that are critical for normal liver physiology and injury response (1). In their normal state within the liver, HSCs are quiescent and help maintain liver structure, regulate extracellular matrix (ECM) turnover, and store vitamin A. However, in response to liver injury, HSCs transition to an activated state through myofibroblast transdifferentiation. This activation triggers HSC proliferation and increased ECM secretion, resulting in scar formation at the injury site(s) and the production of cytokines and growth factors that promote the proliferation of other liver cell types. Prolonged HSC activation can lead to liver fibrosis and cirrhosis. Identifying mechanisms that impact HSC development can offer valuable insights into pathways involved in liver development, pathophysiology, and regeneration.

We and others previously identified *podocalyxin-like* (*podxl*) as a gene whose expression is highly enriched in adult HSCs in zebrafish (2, 3). In humans, *PODXL* belongs to the CD34 gene family that is comprised of *PODXL, CD34*, and *ENDOGLYCAN* (*PODXL2*)(4). It is a transmembrane protein recognized for its role in enhancing cell proliferation, invasion, and migration (5). *Podxl* null mice demonstrate severe defects in renal development accompanied by 100% perinatal mortality (6), and morpholino knockdown of *podxl* in zebrafish leads to disruption of the characteristic kidney podocyte architecture (7), supporting a conserved role for *podxl* in kidney development. However, the role of *podxl* in liver development has not been explored.

In the present study, we sought to define the role of *podxl* in zebrafish liver development using a combination of genetic approaches. This manuscript is based in part upon A.N.R.’s dissertation (8). We found that *podxl* knockdown using CRISPR/Cas9 or an ATG morpholino (7) resulted in a significant decrease in HSCs at 6 days post-fertilization (dpf). In contrast, *podxl* deletion mutants showed either no significant change in HSC number (three mutants) or a significant increase in HSC number (two mutants). *Podxl* mutants injected with *podxl* CRISPR/Cas9 knockdown, unlike wildtype control siblings, did not show a decrease in HSC number, supporting the hypothesis that decreased HSC number in *podxl* CRISPants is due to disruption of the *podxl* gene rather than to non-specific or off-target effects. Quantitative PCR analysis of *podxl* transcripts in adult mutant livers revealed mRNA and pre-mRNA levels similar to those of wildtype control siblings. CRISPR/Cas9 knockdown of the only other zebrafish *podxl* family member, *endoglycan (endo),* did not affect HSC number in wildtype zebrafish or *podxl* mutants. qPCR of *endoglycan* show similar expression levels between *podxl* mutants and wildtype control siblings. RNA sequencing of *podxl* mutants and wildtype control siblings showed no significant change in transcript levels of genes with sequence similarity to *podxl*, but it demonstrated upregulation of a complex network of ECM-related genes in response to *podxl* loss. Together, these data support a role for *podxl* in zebrafish HSC development and suggest that multiple genes are upregulated to compensate for genomic *podxl* loss.

## Results

### Knockdown of *podxl* in zebrafish results in fewer HSCs

As a first step to explore the role of *podxl* in zebrafish liver development, we knocked down *podxl* using CRISPR/Cas9 technology, injecting Cas9 protein and guide RNA targeting the first exon of *podxl* (sgRNA #1, S1 Table) into single-cell embryos (Fig 1). Knocking down genes in this manner using CRISPR/Cas9 (“CRISPant” larvae) typically results in mosaic animals with diverse indel mutations; mutagenesis rates of 75-99% have been reported in CRISPants (9), but some cells will have monoallelic and/or in-frame mutations that may not affect gene function. To visualize HSCs, we performed the knockdown in *Tg(wt1b:eGFP)* transgenic zebrafish that express eGFP in HSCs (10). As a control, we knocked down tyrosinase (*tyr*) in age-matched siblings. The *tyr* gene encodes an important enzyme in the melanin production pathway (11), and *tyr* knockdown is not predicted to impact liver development.

**Figure 1.**
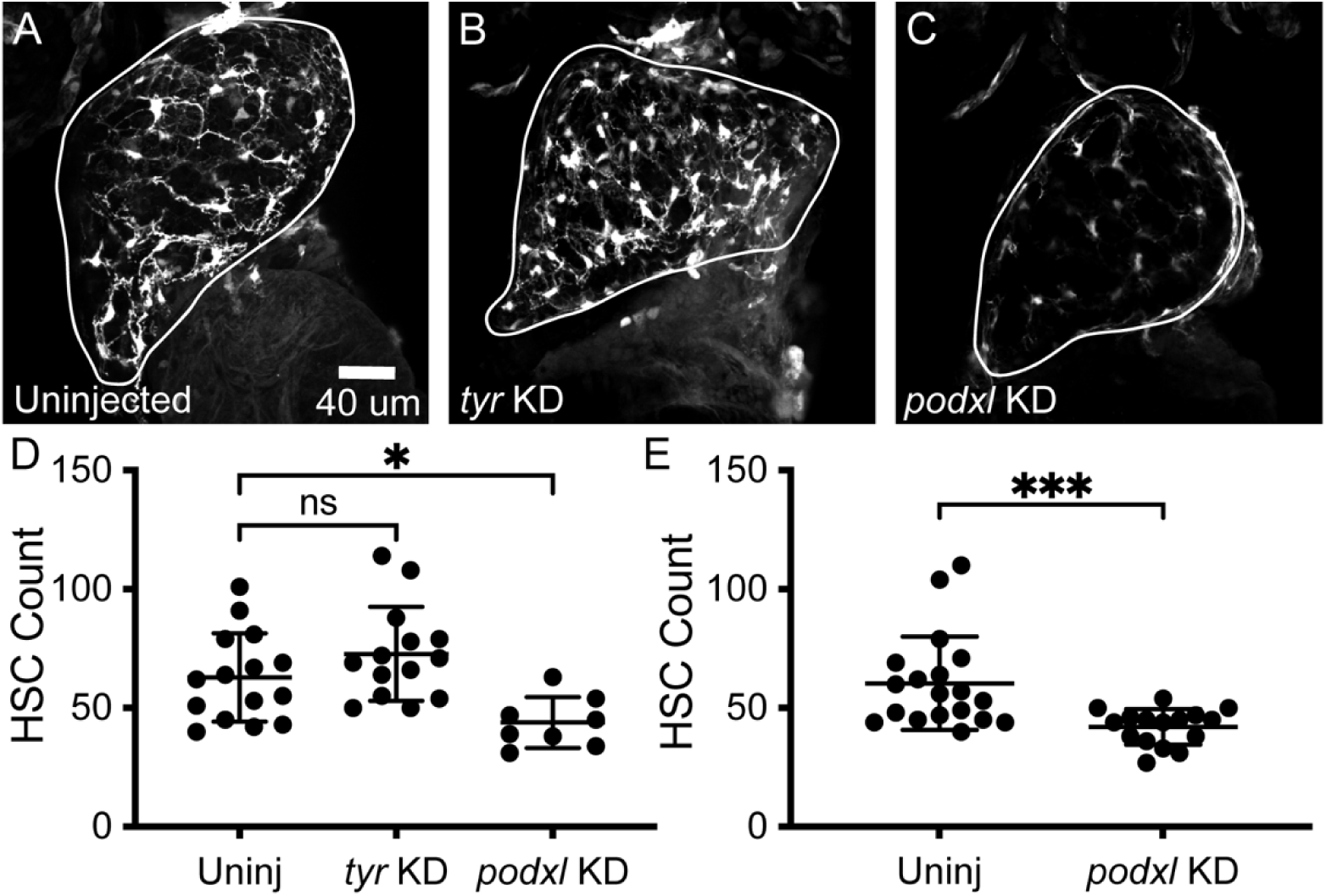
*podxl* knockdown decreases HSC number. *Tg(wt1b:egfp)* zebrafish injected with CRISPR/Cas9 targeting either *tyr* or *podxl*, examined at 6 dpf. Representative confocal projections of uninjected (A), *tyr* KD (B), and *podxl* sgRNA #1 KD (C) (livers outlined in white) and quantified HSC counts (D). (E) HSC counts of 6dpf *podxl* sgRNA #2 KD larvae and uninjected controls. Bars show mean +/- SD. Brown-Forsythe and Welch ANOVA tests (D) and Welch’s t test (E). NS, not significant; *, p<0.05; ***, p<0.001.

We analyzed HSC number at 6 days post-fertilization (dpf) by performing confocal microscopy and manually counting all HSCs in each liver. We found that knockdown of *podxl* using CRISPR/Cas9 (CRISPant larvae) decreased the number of HSCs from 63 ± 19 (mean ± standard deviation) HSCs per liver in uninjected control siblings to 44 ± 11 HSCs per liver in *podxl* CRISPants (31% decrease; p < 0.05; Fig 1A, C, D). Knockdown of *tyr* had no significant impact on the number of HSCs (73 ± 20 HSCs per liver; Fig 1B, D). Knocking down *podxl* in a similar manner but targeting a different exon (exon #2, sgRNA #2, *podxl* KD #2) (S1 Table) decreased the number of HSCs by 30% (60 ± 20 HSCs per liver in uninjected control siblings versus 42 ± 7 HSCs per liver in *podxl* KD#2; p < 0.001; Fig 1E). Successful mutagenesis of *podxl* was confirmed using high-resolution melt analysis (HRMA) (S1 Fig), and *tyr* knockdown was confirmed by identifying decreased pigmentation using brightfield microscopy (S2 Fig).

To determine mutagenesis efficiency in our CRISPant larvae, we examined the *podxl* locus in DNA extracted from larvae that were injected with *podxl* sgRNA #1 and from uninjected control siblings. Using fluorescein-labeled primers and capillary electrophoresis, we determined the percent of allele mutation by calculating the fraction of mutant amplicon product that did not migrate like wildtype amplicon product, suggesting insertions, substitutions, and/or deletions (12). We found that 62% of *podxl* alleles in injected larvae showed mutations compared to 0% of uninjected controls (S3 Fig).

As a complementary approach to determine the effect of *podxl* knockdown on HSC development, we knocked down *podxl* using an ATG morpholino targeting *podxl* (S1 Table) (7). We injected each zebrafish embryo with up to 4.93 ng of morpholino, as using greater than 5 ng has been reported to decrease the specificity of this approach (13). Using 4.93 ng of morpholino, we saw that *podxl* morphants had 47 ± 22 HSCs per liver, compared to 77 ± 21 HSCs per liver in siblings injected with control morpholino (56% decrease; p < 0.001; S4 Fig). With 3.02 ng of morpholino, *podxl* morphants had 46 ± 11 HSCs per liver, compared to 59 ± 16 in uninjected control siblings (23% decrease; p < 0.05; S4 Fig). The finding that multiple methods of *podxl* knockdown produced a similar decrease in HSC number supports the hypothesis that *podxl* is required for normal HSC development.

### The *podxl* CRISPant knockdown phenotype is not due to a developmental delay

We used two approaches to test the possibility that *podxl* knockdown decreased HSC number by delaying development. First, as liver size increases rapidly from 3 dpf to 5 dpf (14), a developmental delay could significantly decrease liver size at 6 dpf. We found that liver size was not significantly impacted by *podxl* CRISPant knockdown (Fig 2A) even when HSC count was significantly reduced (Fig 2B).

**Figure 2.**
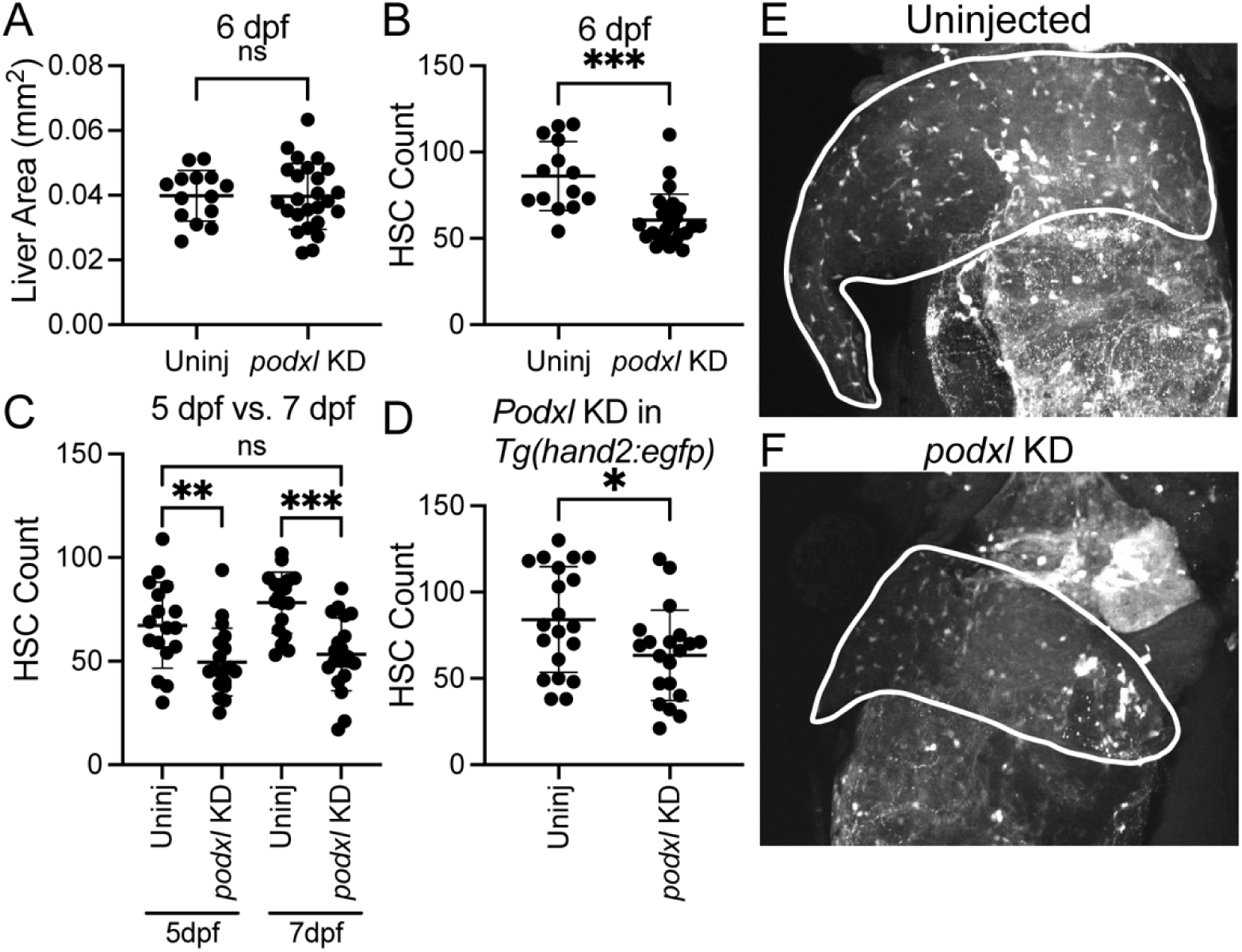
Decrease in HSC number seen with *podxl* KD is not due to developmental delay or an artifact of using the *Tg(wt1b:eGFP)* zebrafish line. Liver area (A) and HSC count (B) of *podxl* KD zebrafish at 6 dpf compared to uninjected siblings. HSC number of *podxl* KD zebrafish at 5 dpf and 7 dpf (C). (D) HSC count of *podxl* KD in *Tg(hand2:EGFP)* zebrafish compared to uninjected siblings at 6 dpf, and representative images (E,F) (livers outlined in white). Bars show mean +/- SD. Welch’s t test (A, B, D) and ANOVA (C). NS, not significant; *,p<0.05; **, p<0.01; ***, p<0.001.

Second, we examined HSC number in *podxl* CRISPant knockdown larvae at different time points. Some genetic manipulations that decrease liver size during larval development result, at least partially, from a transient developmental delay, such that mutant liver size “catches up” to wildtype liver size after hours or days (15). An example of this phenomenon is observed in *wnt2bb* mutant zebrafish, which have very small or absent livers at 50 hours post-fertilization; some animals developing normal functioning livers at later stages (15, 16).

As HSC numbers steadily increase from 3 to 8 dpf (2), a general developmental delay would be expected to decrease the number of HSCs at 6 dpf, rendering it similar to what is seen at earlier time points. If “catch-up” occurs, then differences in HSC number would be less pronounced at later time points. We knocked down *podxl* using sgRNA #1 and examined HSC number at 5 and 7 dpf in age-matched siblings. At 5 dpf we noted 67 ± 21 HSCs per liver in uninjected control larvae and 50 ± 16 HSCs per liver in *podxl* knockdown larvae (25% decrease; p < 0.05; Fig 2C). At 7 dpf we noted 78 ± 15 HSCs per liver in uninjected control larvae and 53 ± 18 HSCs per liver in *podxl* knockdown larvae (32% decrease; p < 0.001; Fig 2C).

The finding that there was not a more pronounced phenotype at 5 dpf than at 7 dpf suggests that *podxl* knockdown does not transiently delay development. This hypothesis is further supported by our finding that 7 dpf *podxl* knockdown larvae still tended to have fewer HSCs than 5 dpf uninjected larvae (p = 0.0502, ANOVA). Taken together, these results argue that there is not a significant developmental delay in the *podxl* CRISPant larvae.

### The *podxl* knockdown phenotype is not specific to the *Tg(wt1b:eGFP)* reporter line

The *Tg(wt1b:eGFP)* transgenic reporter line used to visualize HSCs in the above-mentioned experiments expresses eGFP in HSCs and a few other cell types, including kidney cells (10). The transgenic reporter line *Tg(hand2:eGFP)* can also be used to visualize HSCs, along with lateral plate mesoderm and neural crest cells (2). To test the possibility that the effects of *podxl* knockdown are specific to the *Tg(wt1b:eGFP)* transgenic line, we knocked down *podxl* using *podxl* sgRNA #1 in *Tg(hand2:eGFP)* zebrafish. We found that HSC number at 6 dpf decreased from 84 ± 31 HSCs per liver in uninjected control siblings to 63 ± 26 HSCs per liver in *podxl* CRISPants (25% decrease; p<0.05; Fig 2D-F). This result indicates that *podxl* knockdown decreases HSC number independently of the reporter line used.

### Generation of *podxl* mutants

To further understand the role of *podxl* in the developing zebrafish liver, we generated a series of *podxl* mutants, exploiting the *Tg(wt1b:eGFP)* reporter line to facilitate HSC visualization. Using CRISPR/Cas9 techniques, we made a mutant with a five base pair deletion, resulting in a predicted premature termination codon in the second exon (*podxl ^Ex1,-5bpΔ^*) (Fig 3A, B and S5 Fig). We next made a mutant lacking the entire region between exon 1 and intron 7, leading to a predicted premature stop codon after 18 amino acids (*podxl*^Ex1(p)_Ex7Δ^) (Fig 3C, S6 Fig and S7 Fig).

**Fig 3.**
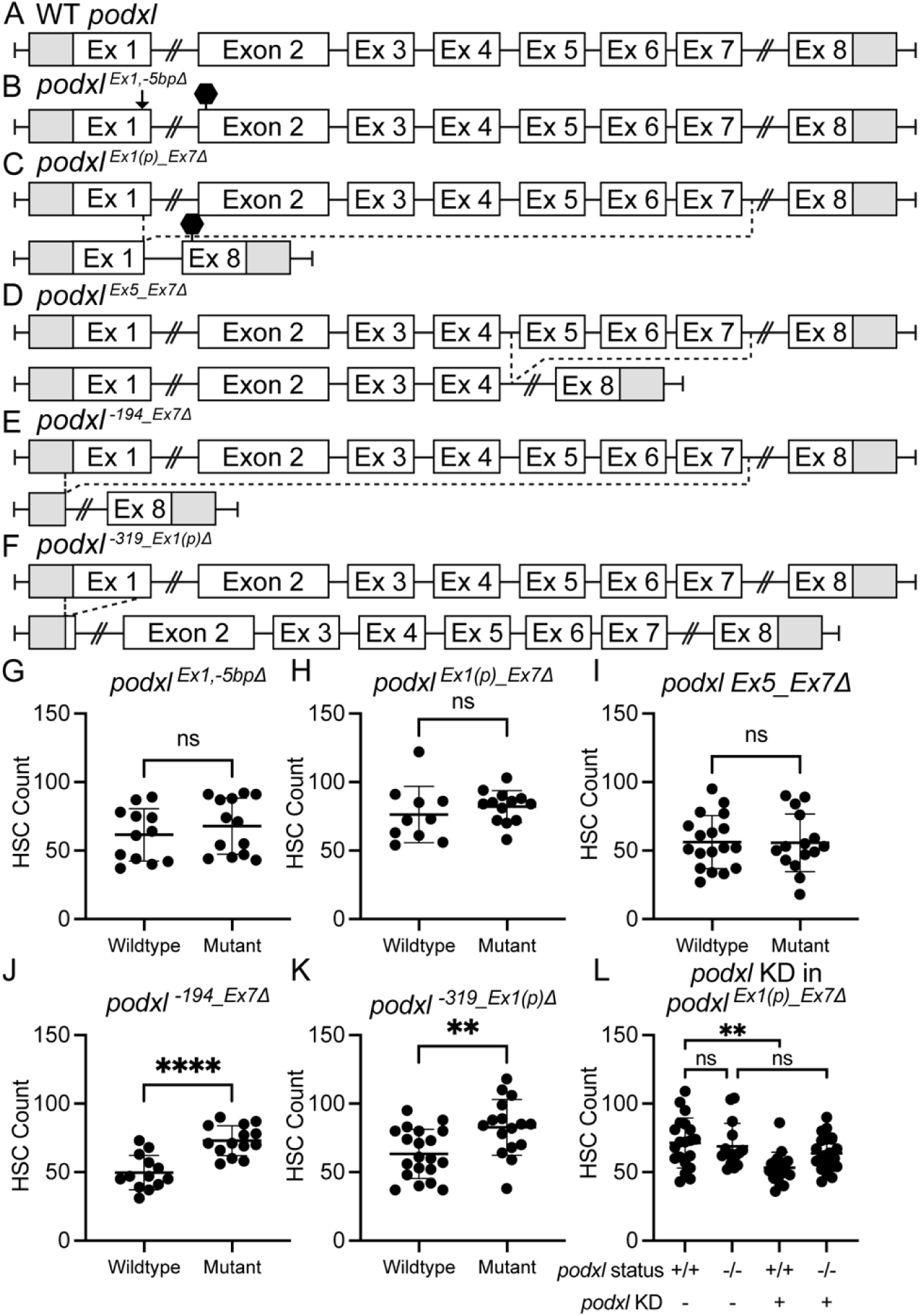
Analysis of HSC number in *podxl* mutant lines shows no change in HSC number for some mutants but an increase in promoter mutants. Schematic representation of the wildtype *podxl* gene (A) and *podxl* mutants (B-F). Deleted regions of *podxl* are depicted by a black arrow (B) or dotted lines (C-F). Premature stop codons are represented by black hexagons (B,C). (G-K) HSC counts of *podxl* mutants compared to wildtype siblings. (L) HSC number in *podxl^Ex1(p)_Ex7Δ^*mutant combined with *podxl* sgRNA #1. Welch’s t-test (G-K). Ordinary one-way ANOVA (L). Bars show mean +/- SD. NS, not significant; **, p<0.01;****, p<0.0001.

Previous experiments examining genetic compensation in mutant zebrafish have shown that premature stop codons can trigger nonsense mediated decay (NMD), which produces degradation products that promote upregulation of gene(s) with similar sequence that compensate for the loss of the gene of interest (17, 18). This form of genetic compensation, termed transcriptional adaptation (17, 19), requires the degradation of mutant mRNA containing a premature termination codon (17, 20). To circumvent the possibility of genetic compensation induced by transcriptional adaptation, we made three additional *podxl* mutants. First, we made an in-frame mutant removing the region between introns 4 and 7 (*podxl*^Ex5_Ex7Δ^) (Fig 3D and S8 Fig), corresponding to the transmembrane domain, which is critical for *podxl* localization and function (21, 22). The *podxl*^Ex5_Ex7Δ^ mRNA lacks a premature stop codon, so it is not predicted to trigger NMD and transcriptional adaptation. We also made two *podxl* mutants lacking the major transcriptional start site marked by H3K4me3 in several tissues including the liver (S9 Fig): *podxl*^-194_Ex7Δ^ (Fig 3E, S10 Fig) and *podxl*^-319_Ex1(p)Δ^ (Fig 3F, S11 Fig). We predicted little *podxl* mRNA would be transcribed in these mutants, making NMD and transcriptional adaptation unlikely.

### Analysis of HSC number in *podxl* mutants

We analyzed HSC number in *podxl* mutants by incrossing *podxl^+/-^*heterozygotes and examining HSC number in *podxl^-/-^* mutants and wild-type control siblings at 6 dpf. We observed no significant difference in HSC number between *podxl ^Ex1,-5bpΔ^* mutants (68 ± 20) and wildtype siblings (62 ± 19) (Fig 3G), *podxl*^Ex1(p)_Ex7Δ^ mutants (82 ± 12) and wildtype siblings (76 ± 21) (Fig 3H), and *podxl*^Ex5_Ex7Δ^ mutants (56 ± 21) and wildtype siblings (56 ± 19) (Fig 3I)

Conversely, in the two promoter mutants, we observed a significant increase in the number of HSCs per liver. We observed a 46% increase in HSC number in the *podxl*^-194_Ex7Δ^ mutant (73 ± 11) compared to wildtype siblings (50 ± 13) (Fig 3J). In the *podxl*^-319_Ex1(p)Δ^ mutant we observed a 32% increase in HSC number (83 ± 20) compared to wildtype siblings (63 ± 18) (Fig 3K).

### *Podxl* mutants are resistant to *podxl* CRISPant knockdown

The discrepant phenotypes in *podxl* knockdown larvae and *podxl* mutants raise two sets of possibilities. First, *podxl* is not important for HSC development, and the *podxl* knockdown phenotype is due to effects on another gene(s). Alternatively, *podxl* is important for HSC development, so acute *podxl* loss causes a decrease in HSCs, but other gene(s) compensate in the case of constitutive/germline *podxl* loss. We would expect that in the first scenario *podxl* CRISPR/Cas9 knockdown would decrease HSC number even in the absence of wildtype *podxl,* while in the second scenario *podxl* mutants would be resistant to *podxl* knockdown.

We distinguished between these two possibilities by knocking down *podxl* in a *podxl*^Ex1(p)_Ex7Δ^ heterozygous incross in the *Tg(wt1b:eGFP)* background. We chose this mutant for these experiments because it lacks the complete cut site for *podxl* sgRNA #1 (S12 Fig). We analyzed HSC number in *podxl*^-/-^ and *podxl*^+/+^ 6-dpf larvae with and without *podxl* sgRNA #1 injection.

Consistent with our prior findings, the HSC numbers of *podxl*^+/+^ larvae were significantly decreased in the group injected with *podxl* sgRNA compared to the uninjected controls. In contrast, we found no significant change in HSC number between the *podxl*^-/-^ groups with and without *podxl* KD injection (Fig 3L). These data support the hypothesis that *podxl* CRISPR/Cas9 knockdown requires an intact *podxl* gene to exert its effects on HSC development.

### Maternal deposition of *podxl* does not compensate for *podxl* loss

Maternal expression and mRNA deposition of certain genes is sufficient to compensate for zygotic loss (23, 24). To address the possibility that maternally deposited mRNAs might compensate for *podxl* loss in mutants, we examined the mRNA-Seq models in the zebrafish ensemble database looking at both pre-zygotic and post-zygotic tracks. We found that *podxl* is not maternally expressed and expression is first detected at 75% epiboly. Additionally, we examined progeny from a *podxl*^Ex1(p)_Ex7Δ^ mutant incross. Homozygous mutant progeny showed no obvious developmental defects and had unremarkable HSC number and morphology (S13 Fig). Homozygous mutant progeny from the *podxl*^Ex1(p)_Ex7Δ^ mutant incross survived to adulthood and were fertile. These data suggest that normal HSC numbers are maintained in *podxl* mutants via a mechanism other than maternally deposited *podxl* mRNA.

### Analysis of *podxl* transcript levels in *podxl* mutants

In some zebrafish mutants, transcriptional adaptation leads to upregulation of related genes, compensating for loss of the gene of interest and blunting the loss-of-function phenotype observed with transient gene knockdown adaptation (17, 19). When transcriptional adaptation occurs, pre-mRNA levels of the gene are close to normal, but mRNA levels are low.

To determine if transcriptional adaptation could be happening in *podxl* mutants, we performed qPCR to examine *podxl* mRNA and pre-mRNA in *podxl* mutant livers alongside wildtype control siblings. We assessed three different parts of the *podxl* mRNA: 5’ UTR, exon 7 to exon 8, and exon 8 (Fig 4, S14 Fig). Additionally, we performed a qPCR on the following sections of *podxl* pre-mRNA: exon 2 to intron 2, intron 2 to exon 3, and intron 7 to exon 8 (Fig 5, S15 Fig). Since some mutants lacked large portions of the *podxl* gene, using three different primer sets for each gene insured that at least one of the sets was predicted to have intact binding sites within each *podxl* mutant transcript (S2 Table).

**Fig 4.**
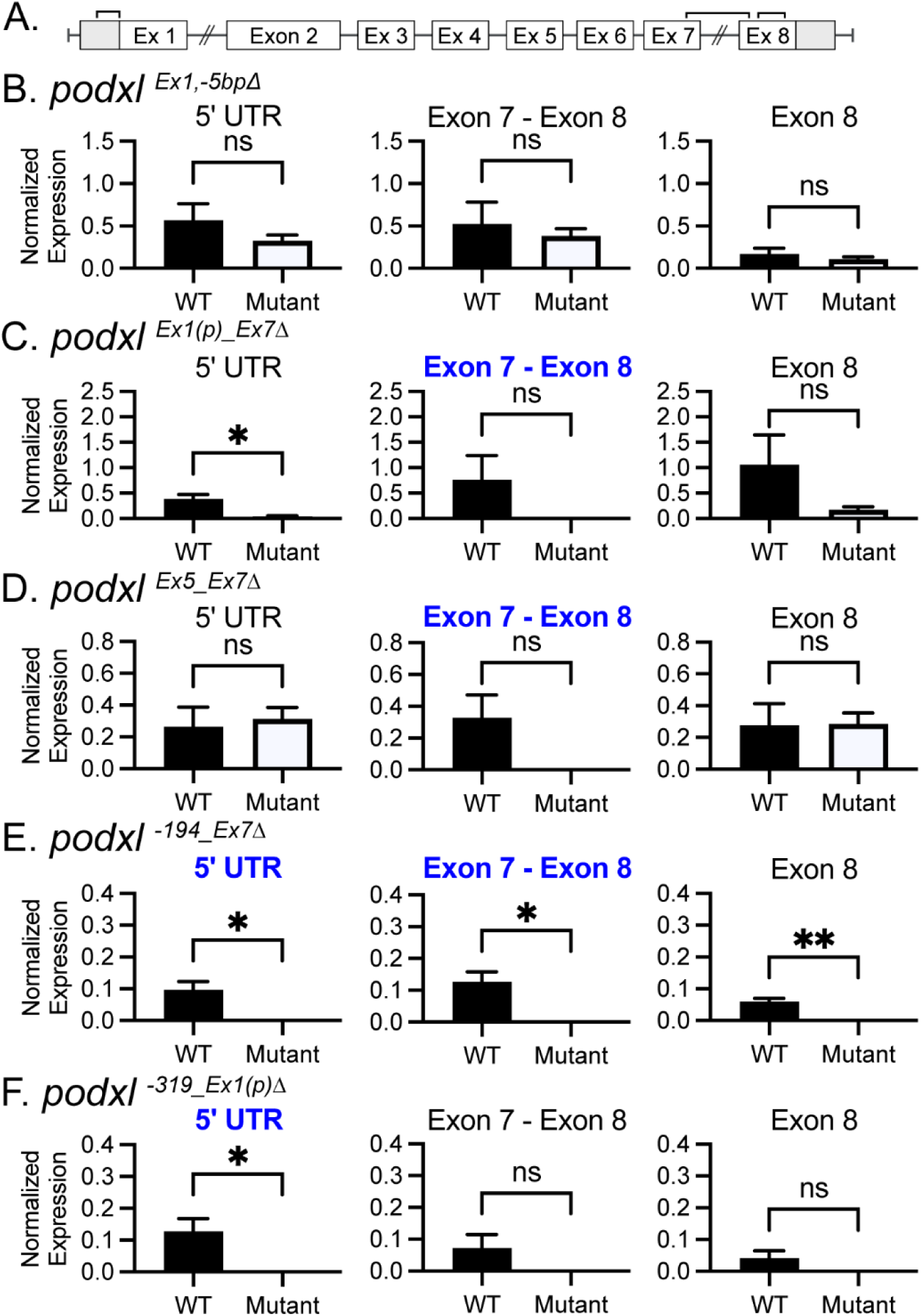
Expression of *podxl* mRNA in *podxl* mutants. (A) Wildtype *podxl* gene schematic with brackets indicating the regions of mRNA amplified by qPCR; 5’UTR - exon 1(left), exon 7 - exon 8 (center), and partial exon 8 (right). Normalized expression levels of each region from the livers of 3mpf *podxl ^Ex1,-5bpΔ^* (B), *podxl*^Ex1(p)_Ex7Δ^ (C), *podxl*^Ex5_Ex7Δ^ (D), *podxl*^-194_Ex7Δ^ (E), *podxl*^-319_Ex1(p)Δ^ (F) mutants and wildtype siblings. Each bar represents the mean +/- SD of three biological replicates (distinct zebrafish). Primers lacking binding sites in the mutant allele are indicated by bold blue graph titles. Welch’s t-test. NS, not significant; *, p<0.05; **, p<0.01.

**Fig 5.**
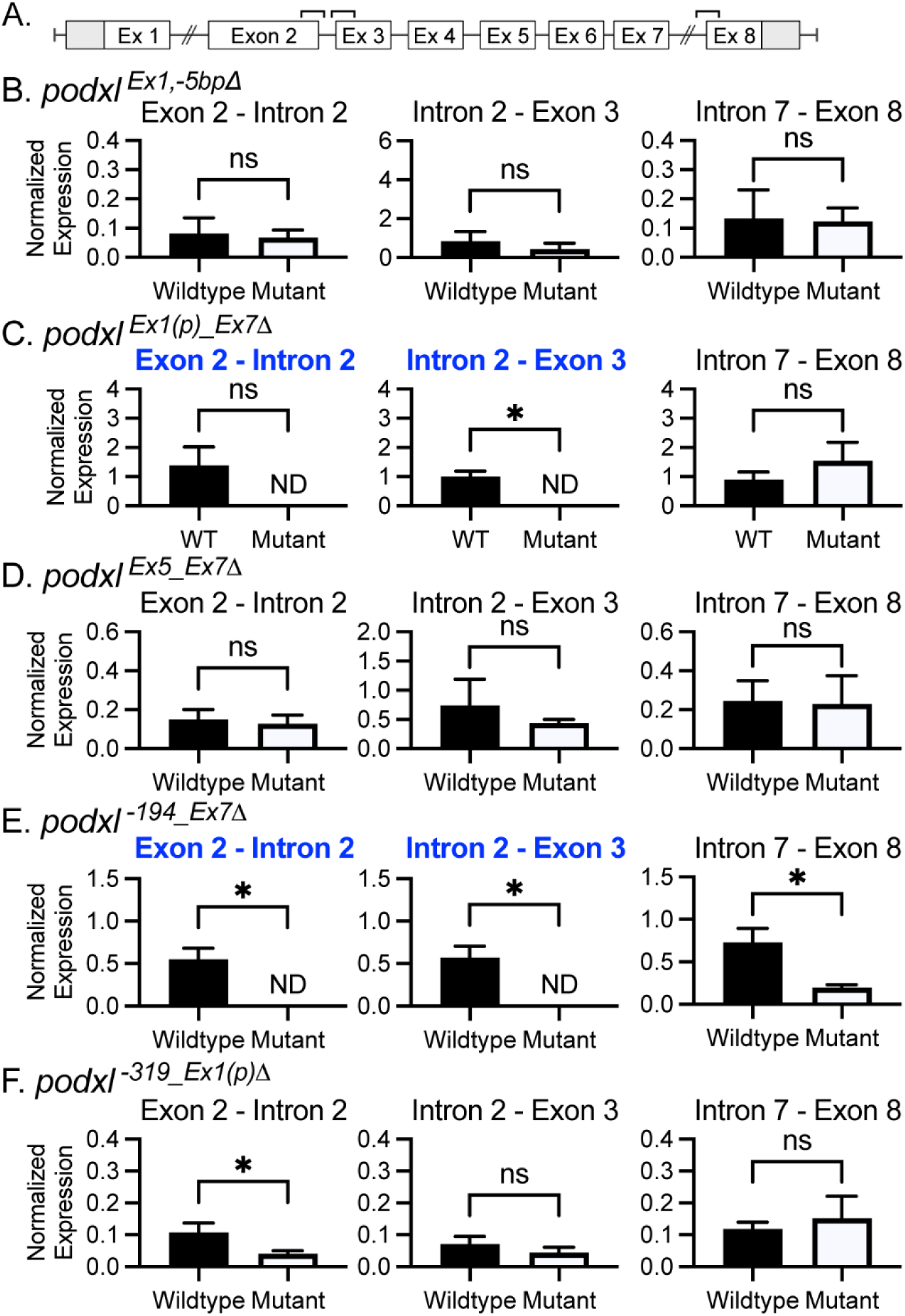
Expression of *podxl* pre-mRNA in *podxl* mutants. (A) Wildtype *podxl* gene schematic with brackets indicating the regions of pre-mRNA amplified by qPCR; exon 2 - intron 2 (left), intron 2 - exon 3 (center), and intron 7 - exon 8 (right).). Normalized expression levels of each region from the livers of 3mpf *podxl ^Ex1,-5bpΔ^*(B), *podxl*^Ex1(p)_Ex7Δ^ (C), *podxl*^Ex5_Ex7Δ^ (D), *podxl*^-194_Ex7Δ^ (E), *podxl*^-319_Ex1(p)Δ^ (F) mutants and wildtype siblings. Each bar represents the mean +/- SD of three biological replicates (distinct zebrafish). Primers lacking binding sites in the mutant allele are indicated by bold blue graph titles. Welch’s t test. NS, not significant; *, p<0.05; ND, not detected.

We found that *podxl* mRNA levels were not significantly different in *podxl ^Ex1,-5bpΔ^* and *podxl*^Ex5_Ex7Δ^ mutants compared to wildtype sibling controls (Fig 4B and 4D). *Podxl* pre-mRNA levels were also not significantly different in these two mutants compared to controls (Fig 5B and 5D). These data suggest that *podxl ^Ex1,-5bpΔ^*and *podxl*^Ex5_Ex7Δ^ mutant pre-mRNA is transcribed normally and *podxl ^Ex1,-5bpΔ^*and *podxl*^Ex5_Ex7Δ^ mutant mRNA is not unstable. On the other hand, *podxl* mRNA levels were significantly reduced in *podxl*^Ex1(p)_Ex7Δ^ mutants (Fig 4C), supporting the hypothesis that mRNA transcription and/or mRNA stability may be decreased in these animals.

The promoter mutants *podxl*^-194_Ex7Δ^ and *podxl*^-319_Ex1(p)Δ^ showed very little *podxl* mRNA expression (Fig 4E and 4F), including the exon 8 region where the qPCR promoters are present in the mutants. The pre-mRNA levels of both promoter mutants were also significantly lower than those observed in wildtype control siblings (Fig 5E and 5F). This result supports the hypothesis that *podxl* transcription is reduced in *podxl*^-194_Ex7Δ^ and *podxl*^-319_Ex1(p)Δ^ mutants; *podxl*^-194_Ex7Δ^ and *podxl*^-319_Ex1(p)Δ^ mRNA may also be unstable.

Although at least two *podxl* mutants (*podxl*^Ex1(p)_Ex7Δ^ and *podxl*^-194_Ex7Δ^) showed a significant reduction in *podxl* mRNA levels, at least some of this decrease might be related to decreased *podxl* transcription, given the decreased *podxl* pre-mRNA levels in *podxl*^-194_Ex7Δ^ and *podxl*^-319_Ex1(p)Δ^. The finding that two *podxl* mutants (*podxl ^Ex1,-5bpΔ^* and *podxl*^Ex5_Ex7Δ^) had normal *podxl* mRNA levels argues that transcriptional adaptation triggered by NMD is not the sole mechanism by which *podxl* mutants compensate for *podxl* loss.

### Upregulation of *endoglycan* does not compensate for *podxl* loss

In some cases of genetic compensation, genes in the same family compensate for the loss of one family member (20). In humans and mice, *PODXL* is part of the *CD34* gene family that includes *PODXL*, *CD34*, and *ENDOGLYCAN* (4). Zebrafish have two CD34 family members: *podxl* and *endoglycan* (*endo*). To determine if *endo* compensates for *podxl* loss, we knocked it down using CRISPR/Cas9 in a *podxl*^Ex1(p)_Ex7Δ^ heterozygous incross (S1 Table). We confirmed cutting of *endo* using HRMA (S16 Fig). We found that there was no significant change in HSC number between the *endo* sgRNA injected *podxl* mutants and the uninjected control mutants (Fig 6A). This result suggests that *endo* is not compensating for the loss of *podxl* in *podxl* mutants.

**Fig 6.**
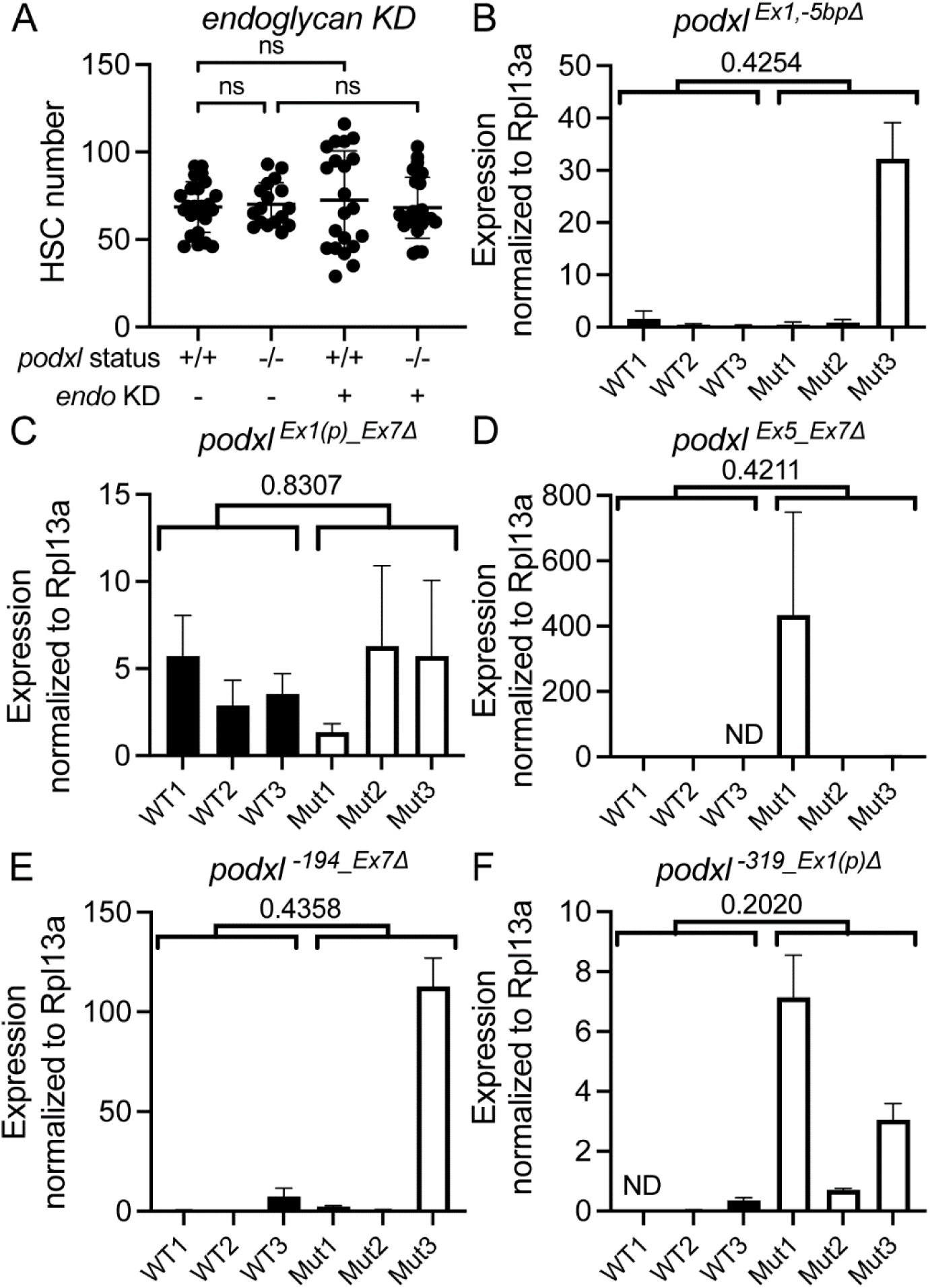
*Endoglycan* does not compensate for *podxl* loss. (A) 6dpf of HSC count in *podxl^Ex1(p)_Ex7Δ^*mutant with CRISPR knockdown of *endoglycan*. qPCR of *endoglycan* in *podxl ^Ex1,-5bpΔ^* (B), *podxl*^Ex1(p)_Ex7Δ^ (C), *podxl*^Ex5_Ex7Δ^ (D), *podxl*^-194_Ex7Δ^ (E), *podxl*^-319_Ex1(p)Δ^ (F). Brown-Forsythe and Welch ANOVA test (A), Welch’s t test (B-F). Bars show mean +/- SD. ns, not significant; ND, not detected.

To further test the possibility that *endo* might compensate for *podxl* loss, we performed qPCR for *endo* on *podxl* mutant livers (S2 Table). Though we occasionally observed increased *endo* expression in some *podxl* mutants, overall there was no significant difference in *endo* expression between *podxl* mutants and wildtype sibling controls (Fig. 6B-F).

### Response to liver injury is unaffected in *podxl* mutants

Some mutants appear unremarkable under normal physiologic conditions, but a phenotype is revealed in response to stress (25). To determine if *podxl* mutants might respond differently to liver injury, we subjected *podxl^Ex1(p)_Ex7Δ^* and *podxl^-194_Ex7Δ^* zebrafish larvae to incubation with 2% ethanol or to hepatocyte ablation with metronidazole/nitroreductase (26). In wildtype zebrafish, ethanol treatment increased HSC number as previously reported (2)(S18 Fig). Hepatocyte ablation decreased liver size as previously reported (26), and it also increased HSC density (S19 Fig). We found that HSC numbers and HSC density in *podxl ^Ex1(p)_Ex7Δ^*and *podxl ^-194_Ex7Δ^* mutants were similar to those of wildtype zebrafish in response to ethanol or hepatocyte ablation (S16 and S17 Fig), supporting the hypothesis that *podxl* is not required in HSCs in these contexts.

### A network of extracellular matrix genes is upregulated in response to *podxl* loss

To characterize transcriptional changes in *podxl* mutant livers, we performed RNA-sequencing on adult livers dissected from three *podxl*^Ex1(p)_Ex7Δ^ mutants and wildtype control siblings. We identified 472 genes that were upregulated and 629 genes that were downregulated in *podxl*^Ex1(p)_Ex7Δ^ mutant livers (log2 fold change > 2 and adjusted p-value < 0.05). By performing GO Enrichment Analysis (27), we found that the upregulated genes were enriched for extracellular matrix, extracellular region, and peptidase activity genes (Table 1). Additionally, out of our top 100 most upregulated genes, 15 of those were genes with extracellular regions and three were extracellular matrix genes (S3 Table). Other significantly upregulated genes included *thrombospondin 1a* (*thbs1a*), an extracellular matrix protein (28), and *ezrin a* (*ezra*), a cytoskeleton linker protein that is known to interact with *podxl* (29). Several significantly upregulated genes have been previously reported to be enriched in zebrafish HSCs (2, 3, 30) (S4 Table). Together these data suggest that *podxl* mutants upregulate a network of extracellular matrix genes and other HSC-enriched genes to compensate for the loss of *podxl*.

**Table 1.**
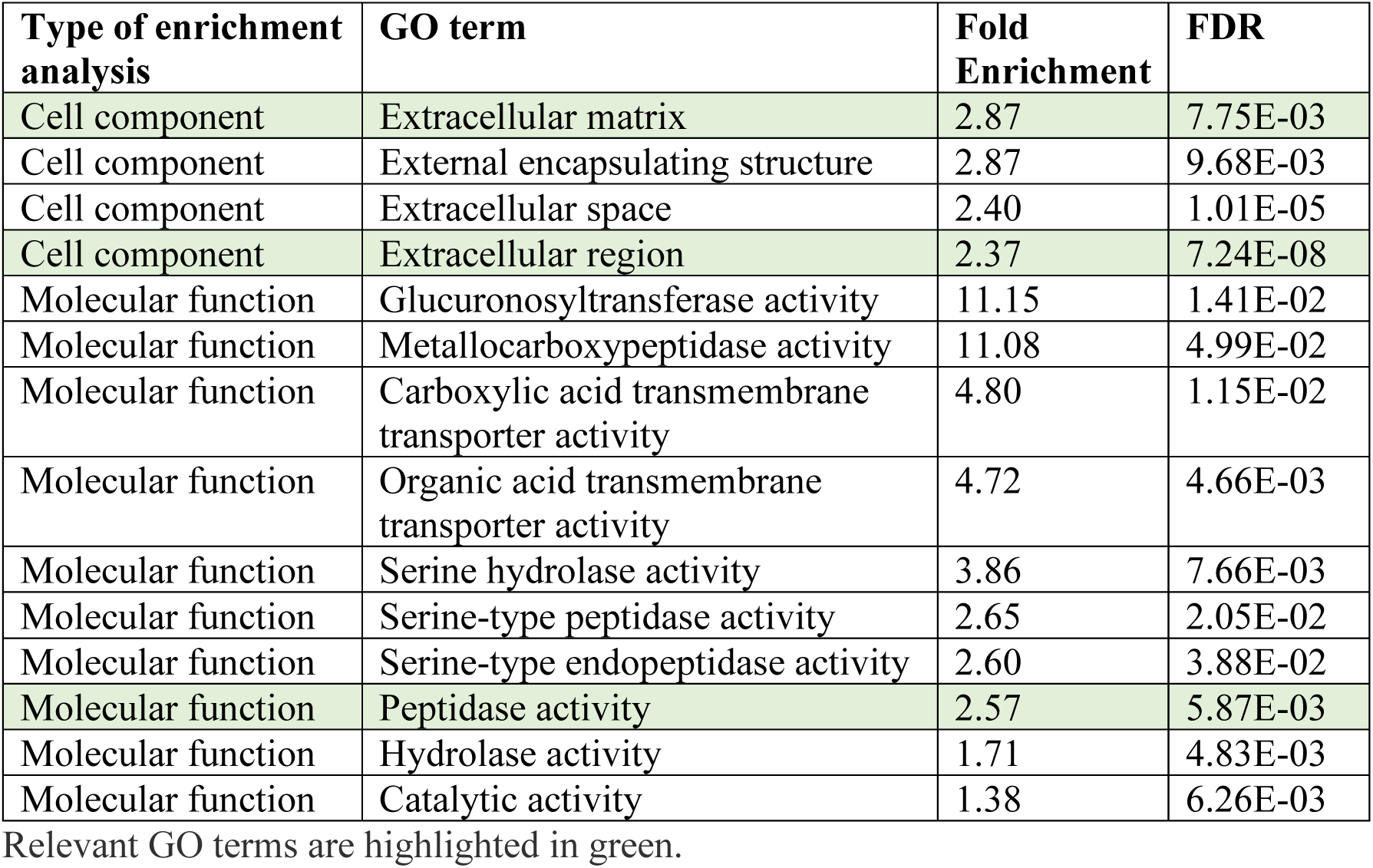
Gene ontology analysis of upregulated genes in *podxl*^Ex1(p)_Ex7Δ^ mutants found using RNA sequencing (27).

We analyzed RNA sequencing data to further investigate the possibility of transcriptional adaptation in *podxl* mutants. If transcriptional adaptation were occurring, genes with similar sequences to *podxl* would be expected to be significantly upregulated in *podxl* mutants (17, 18). We used NCBI BLAST (31) to find genes that had either highly similar (S5 table) or somewhat similar (S6 table) sequences to the zebrafish *podxl* gene and analyzed the expression of these genes in *podxl* mutants compared to wildtype control siblings. We found that none of these genes with sequence similarity to *podxl* were significantly dysregulated, consistent with the hypothesis that transcriptional adaptation is not occurring in *podxl*^Ex1(p)_Ex7Δ^ mutants. Expression of the other zebrafish CD34 gene family member, *endo*, was not significantly different in *podxl*^Ex1(p)_Ex7Δ^ mutants compared to wildtype control siblings, being undetected in both groups. We did not observe a significant enrichment of differentially expressed genes within 700 kbp of the *podxl* start site in *podxl*^Ex1(p)_Ex7Δ^ mutants compared to wildtype control siblings (S7 Table).

We also examined RNA sequencing reads to confirm the *podxl*^Ex1(p)_Ex7Δ^ mutation. We found that *podxl* transcripts in the deleted region were not detected and *podxl* transcripts outside the deleted region were very low (S17 Fig), confirming the presence of the mutation and supporting the qPCR results.

## Discussion

Here we report that *podxl* CRISPant knockdown in zebrafish significantly and robustly decreases HSC number, supporting a role for *podxl* in zebrafish liver development. We found that the decrease in HSC number seen in *podxl* CRISPant zebrafish was not due to a transient developmental delay or to an artifact of the transgenic reporter line. As *podxl* promotes migration and proliferation in colorectal cancer and spermatogonial stem cells (32, 33), it is possible that loss of *podxl* leads to decreased HSC migration into the liver and/or decreased proliferation of HSCs and/or HSC precursors.

We generated and characterized a series of *podxl* deletion mutants and found they had either no change in HSC number or a significant increase in HSC number, in sharp contrast to what we observed with acute *podxl* knockdown. Several lines of evidence argue that the discrepant phenotypes in *podxl* CRISPant knockdown zebrafish compared to *podxl* mutants are due to genetic compensation in *podxl* mutants rather than to an off-target effect of *podxl* knockdown. First, we noted a similar decrease in HSCs with *podxl* knockdown in eight independent CRISPant experiments using two different sgRNAs and performed by two different lab members. Second, *podxl* ATG-morpholino injection, which is expected to decrease Podxl levels by a distinct mechanism involving disruption of translation (7), also decreased HSC number, though at least some of this decrease may not be specifically due to targeting of *podxl* in the liverThird, *podxl* mutants were resistant to *podxl* knockdown.

Genetic compensation represents a highly conserved process that enables organisms such as fly, yeast, mice, and zebrafish to maintain their fitness despite losing an important gene (19, 34). Three major categories of genetic compensation have been described: redundant genes, protein feedback loops, and transcriptional adaptation (19). Redundant genes are expressed in the same tissues and have overlapping functions with the gene of interest, while protein feedback loops lead to the upregulation of genes in the same pathway in response to decreased protein levels (35, 36, 37). The finding that *endo* knockdown in *podxl* mutants did not impact HSC number argues against genetic compensation through redundant genes. The finding that HSC numbers were decreased in *podxl* knockdown zebrafish, which likely have decreased Podxl levels, suggests that protein feedback loops are not sufficient to compensate for *podxl* loss.

Transcriptional adaptation was first described in zebrafish, where Rossi *et al.* found that loss of the epidermal growth factor-like domain multiple 7 (*egfl7*) gene leads to severe vascular phenotypes in morphants though *egfl7* mutants show no apparent phenotype. Further investigation revealed that this compensation is likely due to the upregulation of members of the *emilin* gene family, which share significant sequence similarity with *egfl7* (17, 18). Like Rossi *et al.,* we found that *podxl* mutants were resistant to *podxl* CRISPant knockdown, and three *podxl* mutants showed decreased mRNA levels, compatible with mRNA degradation. On the other hand, two *podxl* mutants showed mRNA levels that were not significantly decreased, suggesting that mRNA degradation is not necessary for genetic compensation in at least some types of *podxl* mutants. Additionally, while Rossi *et al*. found differential expression of only one gene in *egfl7* mutants, our RNA sequencing data showed a plethora of differentially expressed genes in *podxl* mutants. Thus, it seems like genetic compensation in *podxl* mutants occurs through a mechanism that is distinct from the transcription adaptation that occurs in *egfl7* mutants.

We observed an increase in HSC number in two *podxl* mutants, *podxl*^-194_Ex7Δ^ and *podxl*^-319_Ex1(p)Δ^, both of which disrupt exon 1 and the canonical promoter region of *podxl*. Genetic overcompensation has been reported in zebrafish with frameshift mutations in exon 1 of the BMP signaling-promoting gene *marcksb*: compared to wild-type control embryos, *marcksb^ihb199^* embryos exhibit increased Bmp2b fluorescence and increased p-Smad1/5/9 intensity, likely due to upregulation of MARCKS-family members and MARCKS-interacting protein Hsp70.3 (38). It is possible that disruption of the canonical promoter region in *podxl*^-194_Ex7Δ^, *podxl*^-319_Ex1(p)Δ^, and/or *marcksb^ihb199^* mutants leads to creation of an orphan enhancer that promotes transcription of genes that are near the *podxl* locus with respect to three-dimensional chromatin structure. In the human and mouse β-globin family, competition between promoters for a common enhancer, termed the locus control region (LCR), determines the developmental order and compensatory expression of related globin genes (39, 40). To further investigate the possibility that an orphan enhancer mediates the increase in HSC numbers observed in *podxl*^-194_Ex7Δ^ and *podxl*^-319_Ex1(p)Δ^ mutants, it would be interesting to examine the long-range chromatin interactions of the residual promoter proximal DNA in the mutants by Hi-C (41) and/or by using a targeted approach such as 3C (42) aimed at the dysregulated genes we uncovered.

*Egfl7* morphants display severe, highly penetrant vascular phenotypes (18). In contrast, the decrease in HSC number observed in *podxl* CRISPant zebrafish is not binary and instead falls on a spectrum, making statistical analysis necessary to define the phenotype(s) for each experimental group. Thus, although the *podxl* CRISPant phenotype is reproducible, its penetrance is variable, making it challenging to define a compensatory mechanism.

In a recent study of genetic compensation in *slc25a46* mutant zebrafish, researchers found over a hundred dysregulated genes that did not have sequence similarity to *slc25a46* (43). They did not observe mRNA degradation in *slc25a46^238s^* mutants and speculated that the compensatory mechanism in their model may involve a genetic modifier or an orchestrated interaction among multiple genes within relevant networks. It is possible that a similar mechanism is at play in *podxl*^Ex1(p)_Ex7Δ^ mutants, as they also show dysregulation of hundreds of genes. Taken together, the findings in *podxl* and *slc25a46* mutants provide support for a mechanism of genetic compensation that is distinct from redundant genes, protein feedback loops, and transcriptional adaptation.

## Materials and methods

### Zebrafish lines and husbandry

Zebrafish (*Danio rerio*) lines were maintained in compliance with the University of Utah Institutional Animal Care and Use Committee guidelines under standard conditions (44). In addition to male and female wildtype AB strain, the transgenic lines *Tg(wt1b:eGFP)*, *Tg(hand2:eGFP)*, and *Tg(kdrl:mCherry)* were also used. Embryos and larvae were cultured in egg water (2.33 gm Instant Ocean in 1 L Milli-Q water with 0.5 ml methylene blue) and stored in a 28.5°C incubator. Adult zebrafish were maintained on a system of recirculating water and fed powdered food, flakes, and brine shrimp. Animals were euthanized using tricaine methanesulfonate (0.03%) and/or immersion in ice water (rapid chilling).

### SgRNA design and morpholino

Small guide RNAs (sgRNAs) were selected from the top targets identified by CHOPCHOP software (http://chopchop.cbu.uib.no/) with NGG PAM sites and no predicted off-targets. gRNAs were then ordered from IDT and annealed with tracerRNAs at a concentration of 3μM (S1 Table). Morpholino injections were performed consistent with published guidelines (13) with a well-characterized and validated ATG morpholino (7). The morpholino used was ordered from Gene Tools and each embryo was injected with up to 4.93 ng of either standard control or *podxl* morpholino at the single-cell stage.

### CRISPant knockdown experiments

To knockdown *podxl*, *tyr*, and *endo*, we injected single cell embryos with an injection mix that contained 0.25 µg/µL of Cas9 protein, 18 ng/µL *podxl, tyr* or *endo* crRNA, and 33.5 ng/µL tracrRNA. We assessed *podxl* and *endo* knockdown using high resolution melt analysis (HRMA). For *podxl* we used forward primer 5′–GACTGAACGCGGAGAATCTG–3′ and reverse primer 5′– CGTCGTATATGAAGGTAAACTCACC–3′. *Podxl* knockdown experiments were performed by two different lab members to validate the results. For *endo* we used forward primer 5′– GTGCCACGGGATCGAAA–3′ and reverse primer 5′– CCGTCCTCGCTGTCTTC–3′. For *tyr,* gene disruption was assessed by visually inspecting zebrafish larvae for loss of pigmentation.

### Zebrafish embryo genomic DNA extraction

For initial screening and founder screening to check for the presence of *podxl* mutations, ten 2-day old embryos were pooled into a single well. They were digested for 2 hrs at 55 °C in 100 µl DNA digestion buffer (20 mM Tris pH 8, 50 mM KCl, 0.3% IGEPAL, 0.3% Tween20, 0.5 µg/μl Proteinase K). Proteinase K was then inactivated at 95 °C for 10 min. Samples were centrifuged at 2,500 rpm for 2 minutes; 1 µl was used in 12.5 µl PCR reactions.

### Generating *podxl* mutants

We generated five *podxl* mutants using CRISPR/Cas9 technology. The putative *podxl* promoter region and major transcriptional start site were identified by epigenetic feature annotation (Supplementary Methods). We injected single-cell embryos with 0.25 µg/µL of Cas9 protein and 18 ng/µL *podxl* crRNA + 33.5 ng/µL tracrRNA (S6 Table). The injected (F0) zebrafish were screened for germline transmission by crossing with wildtype zebrafish and screening the F1 progeny for mutations using PCR (S9 Table). Temperature cycles were as follows: 98°C for 5′; 35 cycles of 95°C for 30′′, 58-68.3°C (S9 Table) for 30′′, 72°C for 35′′; 72°C for 5′; and 10°C hold. PCR samples were loaded into a 2% agarose gel in TAE.

### Whole-mount Immunofluorescence

Larvae were fixed overnight in cold 4% PFA. After fixation, larvae were rinsed in cold PBS and the skin was removed to reveal the liver (45). To better visualize the eGFP expression in the *Tg(wt1b:eGFP)* and *Tg(hand2:eGFP)* zebrafish, larvae were blocked for at least 1 h with PBS+4% BSA+0.3% Triton X-100 (PBT) and then incubated with chick anti-GFP primary antibody (1:500, Aves Cat #GFP-1020, Lot #0511FP12) for at least 12 h followed by AlexaFluor goat anti-chick 488 secondary antibody (1:200, A11039, Lot #1937504) for at least 12 additional hours.

### Confocal imaging and analysis of hepatic stellate cells

Larvae for all confocal imaging experiments were mounted on their back in 1% low-melt agarose plus SlowFade Diamond Antifade Mountant (Invitrogen) and cover-slipped. Image acquisition for all samples was done using an Olympus FV1000 confocal microscope using the same parameters (HV, gain, offset, etc.) for all zebrafish within an experiment. A 20x lens was used and 3.5-μm thick z stacks were collected imaging the entire liver. Images were blinded and randomized using R (45, 46) and the number of hepatic stellate cells was counted manually from Z-projections of all liver images using Fiji software (ImageJ) (47, 48).

### Larval ethanol exposure and hepatocyte ablation

Two *podxl* mutants were selected for liver injury experiments; *podxl*^-194_Ex7Δ^ and *podxl*^Ex1(p)_Ex7Δ^. Ethanol exposure was performed as previously described (2) on progeny of a *podxl*^+/-^ incross. Larvae were exposed to ethanol at 4 dpf by transferring them to 2% ethanol diluted in egg water. After 24 hours of incubating in ethanol, larvae were moved to fresh egg water at 5 dpf. Larvae were euthanized and fixed in PFA at 6 dpf. After genotyping, wildtype and mutant larvae were dissected to expose the liver and imaged by confocal microscopy.

Hepatocyte ablation was performed with a metronidazole (MTZ)-nitroreductase (NTR) ablation system (26). Larvae from an incross of *podxl*^+/-^ expressing *Tg(wt1b:egfp)*, with one parent expressing *Tg(fabp10:CFP-NTR),* were exposed to either 10mM MTZ or DMSO from 3.5 dpf to 5 dpf. At 6 dpf larvae were euthanized and fixed in PFA. After genotyping, wildtype and mutant larvae were dissected to expose the liver and imaged by confocal microscopy.

### RNA extraction

Total and pre-mRNA was isolated from adult zebrafish livers using the Qiagen RNeasy Mini kit. We euthanized adult zebrafish by ice-water immersion, dissected out their livers, added Qiagen RLT buffer, ground with a pestle, vortexed on ice for 20 minutes, and then followed the Qiagen RNeasy Mini kit protocol. DNAse I was used to remove all traces of DNA from the samples. RNA concentration and purity were quantified using NanoDrop.

### Real-time quantitative PCR

cDNA was synthesized using SS3 Vilo Synthesis kit with OligoDt (Invitrogen) or Maxima First Strand cDNA Synthesis Kit (Thermofisher). Quantitative RT-PCR was performed using PerfeCTa Sybr Green FastMix (QuantaBio, Cat # 95072-012) with the primers listed in S9 Table. The PCR conditions were: 95°C for 3′; 45 cycles of 95°C for 15′′, 60°C for 45′′; melt curve analysis. The fold changes were calculated relative to *Rpl13a* expression (2, 49).

Three technical replicates (three wells) were performed for each sample and were used to generate standard deviations. Melt curves were examined for all wells, and only Ct values with distinct Tm peaks (valid melt curves) were included in the analysis. If one well lacked a valid melt curve, the other two wells were used for the analysis. If two or three wells lacked valid melt curves, that sample was considered not detected. P-values were derived using Welch’s t-test on the normalized mean expression for wildtype and mutant.

### RNA sequencing

Total RNA was isolated from cell lines using DirectZol RNA Miniprep kit according to the manufacturer’s instructions (Zymo). The concentration and purity were quantified using NanoDrop. The quality of the preparation was considered adequate based on the lowest RIN value of 9.3. Library preparation, sequencing, and analysis were performed at the University of Utah High-Throughput Genomics Shared Resource using the Illumina TruSeq Stranded mRNA Library Prep (Illumina) and NovaSeq S4 Flow Cell 2 platform (Illumina) according to the manufacturer’s instructions.

### CRISPR efficiency testing by fragment analysis

Injected and uninjected control larvae were euthanized and DNA was extracted using the protocol stated above. PCR was performed using FAM-labeled primers ordered from IDT (Forward primer 5′–/56-FAM/AGACGAAAAGCGGAACCGAG–3′; Reverse primer 5′–/56-FAM/CCTCGTCGTATATGAAGGTAAACTC–3′) and the KAPA2G HotStart PCR Kit (KAPA Biosystem, #KK5501). The PCR conditions were as follows: initial denaturation step at 95 °C for 3 min followed by 35 cycles at 95 °C for 12 sec (denaturation), 68 °C for 12 sec (extension), and 72 °C for 5 sec (elongation). The final elongation step was 72 °C for 10 min. Samples were analyzed by the University of Utah DNA Sequencing Core Facility.

### Quantifying larval liver size

Larvae were euthanized at 6 dpf using tricaine methanesulfonate (0.03%) and fixed in 4% paraformaldehyde (PFA) overnight at 4 °C with gentle rocking. Larvae were then rinsed out of PFA using cold 1X PBS and the skin surrounding the liver was removed with forceps (45). Larvae were mounted in 3% methyl cellulose and images of the left side of each liver were taken using a dissecting microscope on the lowest magnification. Images were blinded and liver size was quantified using FIJI/ImageJ (47).

### Identifying *podxl* homologs, genes with sequence similarity to *podxl,* and HSC-enriched genes

To identify *podxl* homologs, we searched ensembl genome browser and UCSC genome browser for *cd34* and *endoglycan* (*podxl2*). To look for genes that had sequence similarity to *podxl*, we used NCBI BLAS to query *Danio rerio* Reference RNA Sequences and either highly similar sequences or somewhat similar sequences. We examined all genes with a 5% query cover or higher.

We examined genes that were significantly upregulated in *podxl* mutants (log2fold change> 2, padj<0.05) alongside three publicly available datasets for HSC-enriched genes. Genes meeting criteria for HSC enrichment and/or upregulation in at least two of these four datasets were included in S4 Table. “Mutant” column indicates that the gene was upregulated with a log2fold change>2 and padj<0.05 in adult zebrafish *podxl* mutant livers compared to *podxl* wildtype livers (RNA sequencing experiment reported in this manuscript; data to be deposited in GEO upon manuscript acceptance). “Yin” column indicates that the gene was among the top 500 most upregulated (greatest log2fold change) in FACS-sorted HSCs from adult zebrafish livers compared to non-fluorescing other liver cell types (2). “Morrison” column indicates that the gene was upregulated with a log2fold change>2 and padj<0.05 in endothelial/HSC clusters upon single-cell RNA sequencing of wildtype zebrafish livers (30). “Spanjaard” column indicates that the gene was one of 50 genes denoted as characteristically expressed in HSCs in adult zebrafish (3).

## Acknowledgements

We thank Jerry Kaplan for critical review of the manuscript, Director of Aquatics Carrie Barton and the entire Office of Comparative Medicine aquatic animal care team for outstanding zebrafish care and husbandry, and Liam O’Brien for experimental assistance. We acknowledge the Cell Imaging Core, the Mutation Generation and Detection Core, and the DNA Sequencing Core at the University of Utah for use of equipment, and we thank Xiang Wang for assistance in image acquisition. We acknowledge support from the High-Throughput Genomics and Cancer Bioinformatics Shared Resource and HCI’s National Cancer Institute Cancer Center Support Grant P30CA042014.

This work was supported by the American Cancer Society (RSG-22-014-01-CCB), the National Cancer Institute (R01CA222570-NIH-NCI), and the Damon Runyon Cancer Research Foundation (DR 61-20).

## Author contributions

A.N.R.: Data acquisition, analysis, and interpretation; N.M.M.: Data acquisition, analysis, and interpretation; S.M.K.: Data acquisition and analysis; J.H.: Data acquisition and analysis; M.T.: Data acquisition; A.S.: Data acquisition; S.A.B.: Data analysis and interpretation; K.J.E.: Conception, experimental design, funding, and data interpretation. The initial manuscript was drafted by A.N.R. and N.M.M. and reviewed critically for important intellectual content by all other authors.

