## Supporting Information for "Genetic compensation in *podocalyxin-like* mutants during zebrafish liver development"

#### Supplementary Methods

##### Epigenetic feature annotation near *podxl*

Chromatin Immunoprecipitation and sequencing (ChIP-Seq) data from Yang et al. (Yang et al. 2020) were downloaded from GEO (GSE134055). The H3K4me3 and H3K27ac peaks for each tissue in the region from chromosome 4: 11,728,308-11,757,188 relative to zv10(GRCz10), which corresponds to +/- 5kb beyond the *podxl* gene body, were extracted and plotted in R 4.2.2 using the *Gviz* package. The gene model for *podxl* was downloaded from the UCSC Genome Table Browser using the UcscTrack() function from *Gviz* from the danRer10 table with track="NCBI RefSeq" and table= "refGene". The transcript data was collected using the same function with track="Ensembl Genes".

### Supplementary Figures

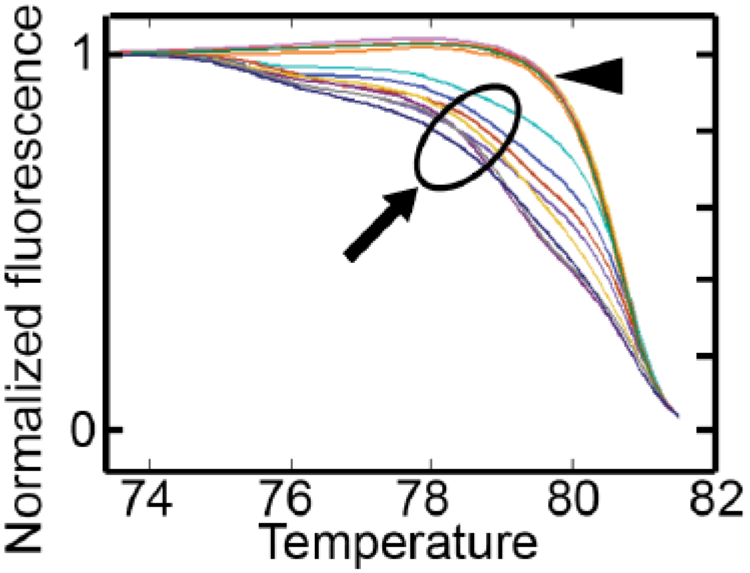


**S1 Fig. High resolution melt analysis of *podxl* knockdown.** Representative shifted melting curves. The presence of *podxl* mutations in every injected embryo was confirmed by amplifying the region surrounding the CRISPR target site and performing high-resolution melt analysis (HRMA). Wild-type *podxl* (arrowhead) and mutated *podxl* (circled, arrow) show distinct shifted melting curves. Only zebrafish with confirmed mutations were included in data analysis.

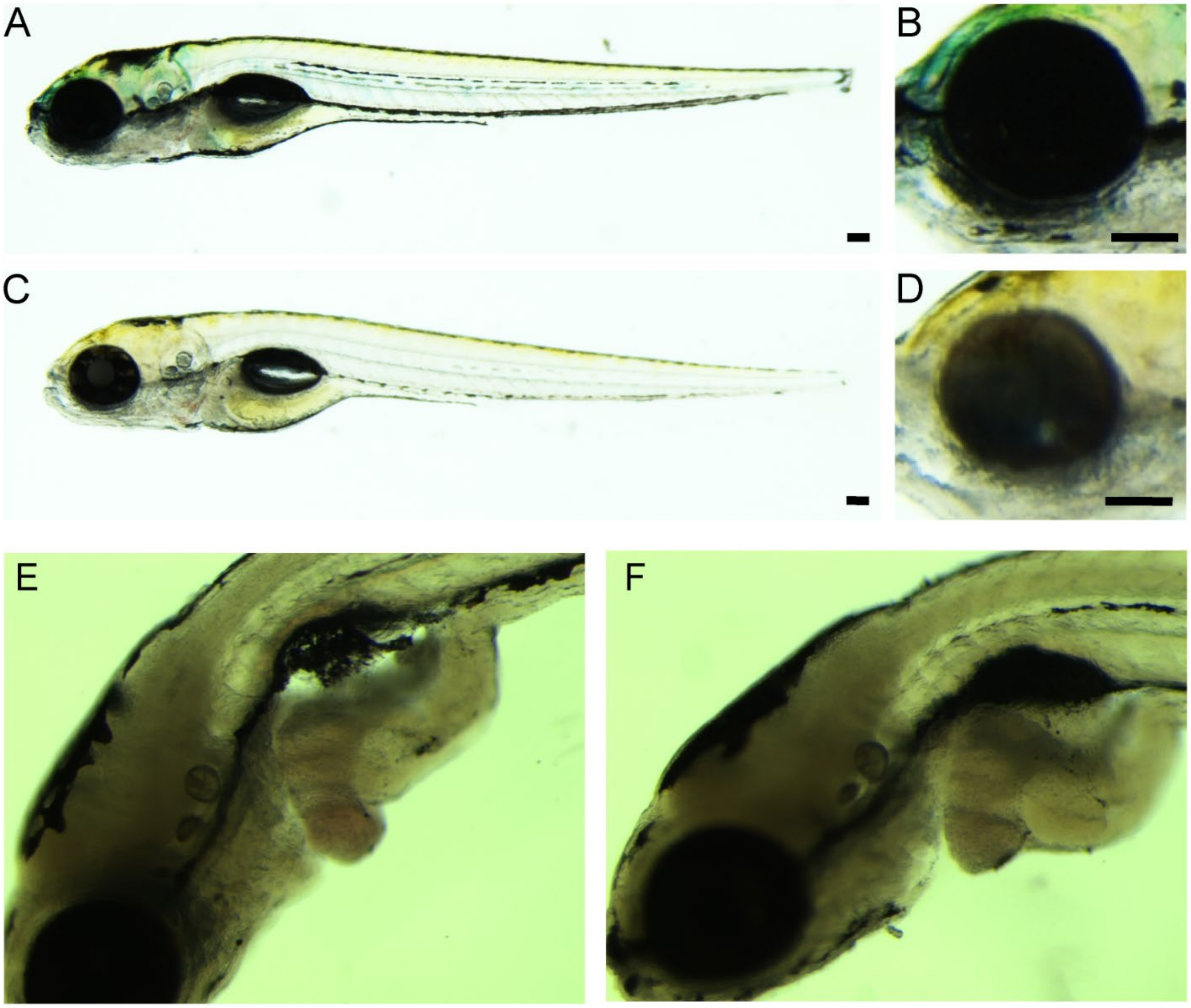


**S2 Fig. Effects of CRISPant injection on morphology by brightfield analysis.** The efficacy of *tyr* knockdown was confirmed by examining zebrafish under brightfield microscopy. Compared to uninjected sibling controls (A and B), zebrafish with successful targeting of *tyr* (C and D) showed overall loss of pigment along the length of the body (A, C) and the eyes (B, D). Scale bars 100 µm. Compared to uninjected sibling controls (E), zebrafish with successful targeting of *podxl* (F) showed no change in liver size or other gross developmental defects.

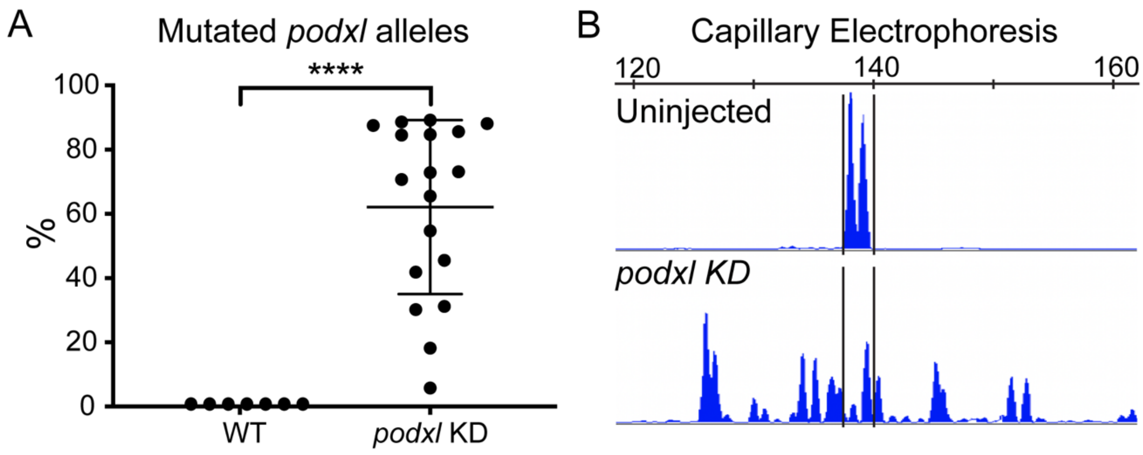


**S3 Fig. Capillary electrophoresis.** (A) The percentage of mutated *podxl* alleles was quantified in individual embryos by capillary electrophoresis. (B) Representative capillary electrophoresis tracings of amplicons from uninjected (top) and injected (bottom) larvae. Welch's t test (A). Bars show mean +/- SD. ****, p<0.0001.

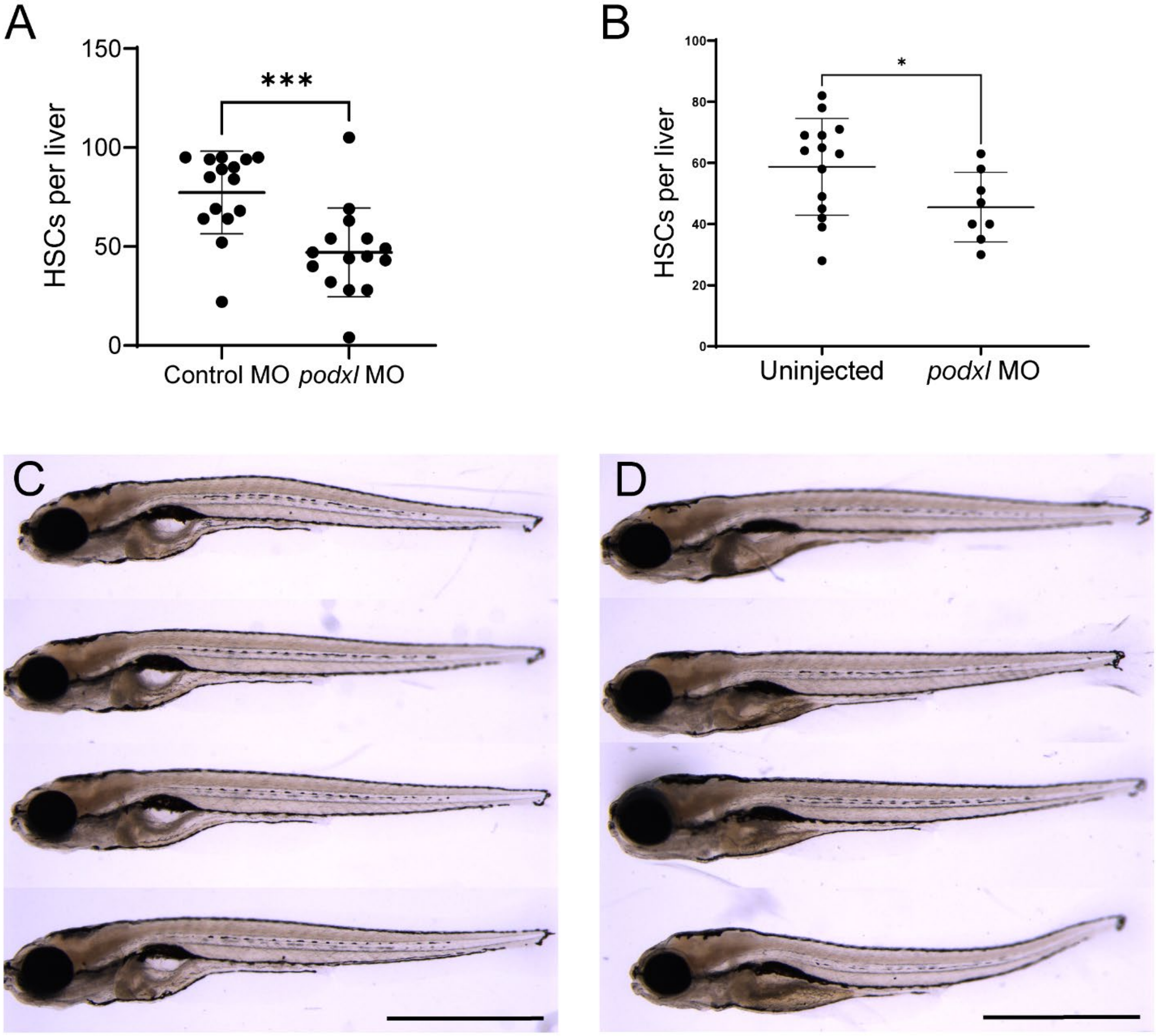


**S4 Fig. Knockdown of *podxl* using a morpholino.** Zebrafish embryos were injected at the one-cell stage with 4.93 ng (A) or 3.02 ng (B) of morpholino targeting *podxl* (S1 table). (A-B) HSCs were examined at 6 days post-fertilization using the *Tg(wt1b:eGFP)* reporter line and confocal microscopy. Statistical analysis was performed using GraphPad Prism (Mann-Whitney test). Bars show mean +/- SD. *, p<0.05; ***, p<0.001. The experiment with 4.93 ng of *podxl* MO (A) was performed twice with similar results. (C-D) Representative brightfield images of uninjected (C) or *podxl* MO-injected (D) larvae from the same injection day and same clutch of larvae that were examined by confocal microscopy and quantified in (B).

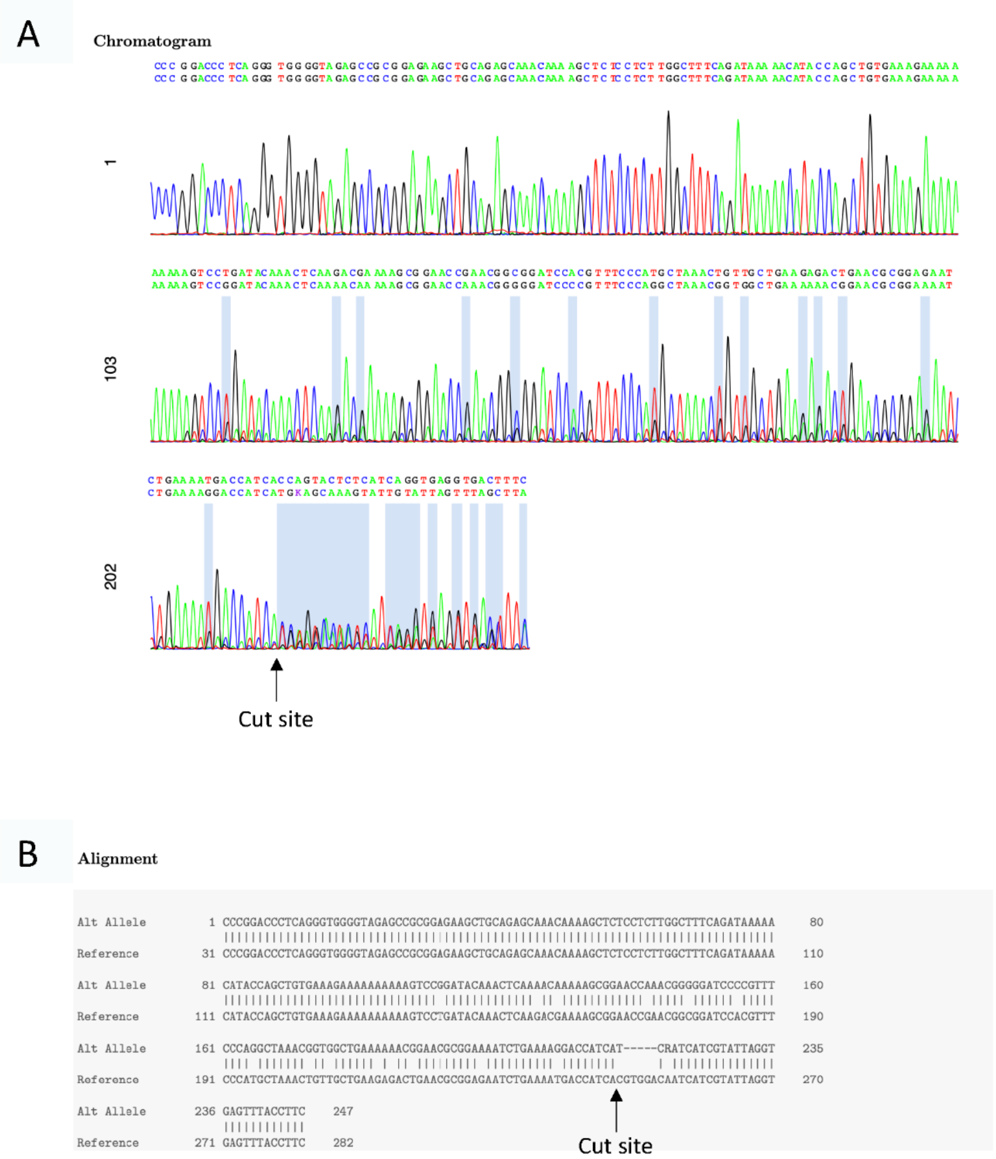


**S5 Fig. Identification of *podxl* $^{Ex1,-5bp\Delta}$ mutant.** We sequenced *podxl* mutant DNA using Sanger sequencing and determined the sequence using Poly Peak Parser (Hill et al. 2014). (A) Chromatogram showing sequence of zebrafish that is heterozygous for the *podxl* $^{Ex1,-5bp\Delta}$ mutation. (B) Alignment of *podxl* $^{Ex1,-5bp\Delta}$ sequence with wildtype sequence.

```
podxl-MUTANT        CAACGCTTCA CAATCCGCGC AGCCAAGGTA ACACACCAAA ACTTTGCCCC
podxl-wildtype      CAACGCTTCA CAATCCGCGC AGCCAAGGTA ACACACCAAA ACTTTGCCCC


podxl-MUTANT        CCAAATCCCC GGACCCTCAG GGTGGGGTAG AGCCGCGGAG AAGCTGCAGA
podxl-wildtype      CCAAATCCCC GGACCCTCAG GGTGGGGTAG AGCCGCGGAG AAGCTGCAGA


podxl-MUTANT        GCAAACAAAA GCTCTCCTCT TGGCTTTCAG ATAAAAACAT ACCAGCTGTG
podxl-wildtype      GCAAACAAAA GCTCTCCTCT TGGCTTTCAG ATAAAAACAT ACCAGCTGTG


podxl-MUTANT        AAAGAAAAAA AAAGTCCTGA TACAAACTCA AGACGAAAAG CGGAACCGAG
podxl-wildtype      AAAGAAAAAA AAAGTCCTGA TACAAACTCA AGACGAAAAG CGGAACCGAG


podxl-MUTANT        CGGCGGATCC ACGTTTCCCA TGCTAAACTG TTGCTGAAGA GACTGAACGC
podxl-wildtype      CGGCGGATCC ACGTTTCCCA TGCTAAACTG TTGCTGAAGA GACTGAACGC


podxl-MUTANT        GGAGAATCTG AAAATGACCA TCA....... .......... ..........
podxl-wildtype      GGAGAATCTG AAAATGACCA TCACGTGGAC AATCATCGTA TTAGGTGAGT


podxl-MUTANT        .......... .......... .......... .......... ..........
podxl-wildtype      TTACCTTCAT ATACGACGAG GATGCAGCTT AAATATAAAC TCGATCTCTT


podxl-MUTANT        .......... .......... .......... .......... ..........
podxl-wildtype      CAGTAATTAA ATATAATGTT AGTCGGTTAA TTTGAGGGAC ATGTGCTGGG


podxl-MUTANT        .......... .......... .......... .......... ..........
podxl-wildtype      TTTATATTGA TGATTAAGTT CAATCAAGGT TGTTTGTAGC CTATAACTTG


podxl-MUTANT        .......... .......... .......... .......... ..........
podxl-wildtype      TTTGAATATT TCACATGTGT ATATATGGCG ATATAACGCC ATGATTTAAA


podxl-MUTANT        .......... .......... .......... .......... ..........
podxl-wildtype      CTATCAAGGG TTAACTTTAA TTCAGTTTCT TATGAATTAA AGTAACGCAC


podxl-MUTANT        .......... .......... .......... .......... ..........
podxl-wildtype      ATAAACAGTC AGAAATTCAG TTGTACATAA TTTTTGTGTG ACCAAACACT


podxl-MUTANT        .......... .......... .......... .......... ..........
podxl-wildtype      TGCCAGCTAG TAAAGGAAGC TTATTGAATC TCCCCAAAGC AATTAGGGTT


podxl-MUTANT        .......... .......... .......... .......... ..........
podxl-wildtype      TAATAATGTT TTCAAAACGT TGACTTCATT AGGTTAATTA TGGGATCATT


podxl-MUTANT        .......... .......... .......... .......... ..........
```

```
podxl-wildtype      TATTAATGAA ACGTTTTTAT TTGGACGATC CTGCAGCATC TTTTAATGCT

podxl-MUTANT        .......... .......... .......... .......... ..........
podxl-wildtype      GCTACTTATA GTTTTCTAAC ACTCACAAAT CCTTAAATTC TCTAAAAACA

podxl-MUTANT        .......... .......... .......... .......... ..........
podxl-wildtype      AGTGCAGGAT TTTAAACTGT TTAAATGCTT TTAAAGTATT CAAAACCTCC

podxl-MUTANT        .......... .......... .......... .......... ..........
podxl-wildtype      ACATTGTGAA CTTTTTTTTA GCTTGTTTGT CTAGTTCTTA CTATTATATT

podxl-MUTANT        .......... .......... .......... .......... ..........
podxl-wildtype      AGTGTCTTTT TAATTTCCCA TGAGTGTTTA CAGAGAACAG TGGTTCCCAT

podxl-MUTANT        .......... .......... .......... .......... ..........
podxl-wildtype      GTACAGCACT GAAAAACAGA AGTAATGTTT AATTTAAACT AATTATCACA

podxl-MUTANT        .......... .......... .......... .......... ..........
podxl-wildtype      AGTGCTCACA TTTACATTTG CAAGCTCTCG AGTTACGCAA CTGATTTACC

podxl-MUTANT        .......... .......... .......... .......... ..........
podxl-wildtype      TTCATAGTTG TGGGGATTGT TCACCCAAAT ATGACACTCA TCCTCAAGTG

podxl-MUTANT        .......... .......... .......... .......... ..........
podxl-wildtype      GTTCCAATCC TTTATGAGTT TCTTTATTCT GTTGAACAAT AAAGATGATA

podxl-MUTANT        .......... .......... .......... .......... ..........
podxl-wildtype      TTTTGAAGAA AGCTGAAAAC CTGTAATCAT TGACATCCGA CATAGGAAAA

podxl-MUTANT        .......... .......... .......... .......... ..........
podxl-wildtype      ACAAATATAC TTGCTTTCCT CAAAATAACC TTCTGGTGTT CAGTAGAAGA

podxl-MUTANT        .......... .......... .......... .......... ..........
podxl-wildtype      AAGAAAGTCA ACAAGTTTGA CACAAGTGAA GGGAAATTTC AGTTTGGGGT

podxl-MUTANT        .......... .......... .......... .......... ..........
podxl-wildtype      GAATTATCCG CATAATAACA AAATATAAAT CAATGTCCAT CTGATGCATC

podxl-MUTANT        .......... .......... .......... .......... ..........
podxl-wildtype      AGAAAAAGAA AGTGCATAAG CATGTTGAAA ATAAGTACAA AATCTGTATA

podxl-MUTANT        .......... .......... .......... .......... ..........
podxl-wildtype      ACCCAAAGCC AACTTGAAGT ATACAGGAAT TAGGATGTTT CATGTTGAAG

podxl-MUTANT        .......... .......... .......... .......... ..........
podxl-wildtype      AGGATAAACC AAAAGGCTGA TAAATCTAAC TCTTCTCTTA TAATAACCAT
```

```
podxl-MUTANT        .......... .......... .......... .......... ..........
podxl-wildtype      TGCAGTAATT CATTTCCAAT TTCTCAGCAG CTTATCAGCA ATTACGCTAC

podxl-MUTANT        .......... .......... .......... .......... ..........
podxl-wildtype      TTGTCTGCGT GCATAAAATC AGTTACTACT GAGTTTACAG CACTGATACT

podxl-MUTANT        .......... .......... .......... .......... ..........
podxl-wildtype      GTGTAACAAT GCCATGTTTT AAAACTTGCA ACCGGAAAAA AGATATGTTG

podxl-MUTANT        .......... .......... .......... .......... ..........
podxl-wildtype      AGATTGCAAA CTTTCTTATC TGGTTGTTTT CTGTTCTCCA CAGGTGCCTT

podxl-MUTANT        .......... .......... .......... .......... ..........
podxl-wildtype      CCTACATGAC ATGGCGTATG CACAGTCTGA AACCACAATA CCCACCGCTT

podxl-MUTANT        .......... .......... .......... .......... ..........
podxl-wildtype      CTGAATCACA CAGTACCTCG AATAGAAGTG TGGTCACTGA TGCAACTGTG

podxl-MUTANT        .......... .......... .......... .......... ..........
podxl-wildtype      TCTACTACGG TGCACAAAGC GACTAATGCA CAAACAACTG AAGCACAAAC

podxl-MUTANT        .......... .......... .......... .......... ..........
podxl-wildtype      AACATCCATT CCAGCTGGTC CAACAAGCCA AGCCACACAT CCTCCCACTT

podxl-MUTANT        .......... .......... .......... .......... ..........
podxl-wildtype      CTGCACCAAC TACCCAATCA GATCCTAACA ACAACCAGGG CTCAACAGCT

podxl-MUTANT        .......... .......... .......... .......... ..........
podxl-wildtype      TTACCCAGTA CATCCACCCT ATACATGCAA AATGTCAGTT CTGCTCAAGT

podxl-MUTANT        .......... .......... .......... .......... ..........
podxl-wildtype      AATACCAACT GGAGTTGACC CAAATACTTC ATTCAGTCCA ACTACCCAAT

podxl-MUTANT        .......... .......... .......... .......... ..........
podxl-wildtype      ACACACTAAC TGATAGTCCT AAATCAGTTA CACAACCGCA TTCATCAGAC

podxl-MUTANT        .......... .......... .......... .......... ..........
podxl-wildtype      CAAGCTTCTA CGTCCAATCC AGATAACCTT AATACTCCGA CTAGCACTTC

podxl-MUTANT        .......... .......... .......... .......... ..........
podxl-wildtype      TGAACGAACA ACACAGCGTG TATCAACTGA ACAAACTACT GCATACAGTC

podxl-MUTANT        .......... .......... .......... .......... ..........
podxl-wildtype      CAGCGAAGCA AGACAGGCCT ACCATTTCTG ACCCAGCCAC ACCAGCTAAC

podxl-MUTANT        .......... .......... .......... .......... ..........
podxl-wildtype      TCTACCACAG AACTCATCAC TTCTAAGCCA GACATTACAA CAGACAAAAA
```

```
podxl-MUTANT        .......... .......... .......... .......... ..........
podxl-wildtype      GACAACAACC GGGACCACAA GTAAGATTTC GCTCACATCA GAATGATCTT

podxl-MUTANT        .......... .......... .......... .......... ..........
podxl-wildtype      AATAACAGAA AAAAATTAAA CATTTAATGT ACAAAAAGGC TCATAAAAAC

podxl-MUTANT        .......... .......... .......... .......... ..........
podxl-wildtype      GAGTAAAATA ATTACAATAA TGAAAAGTAA ATTGTTATTT ATTTTTTATG

podxl-MUTANT        .......... .......... .......... .......... ..........
podxl-wildtype      TCCCTTTTCA GCTCAATTAC TCACCACAGC AAAACCAAAC GCTAGAAGTT

podxl-MUTANT        .......... .......... .......... .......... ..........
podxl-wildtype      TTGTTTCTCA AAGTCTTCAG ACCACAATAG TACCGCAGTC AACAACTGCC

podxl-MUTANT        .......... .......... .......... .......... ..........
podxl-wildtype      CCAATCAAAA CATCCCAGCC TTCGGATCTA CGCAAAAATT TCACTCATGT

podxl-MUTANT        .......... .......... .......... .......... ..........
podxl-wildtype      AAATCTCTCA TTCATGTCTT TCTTACATAT CATCATTACA TTCATAATCC

podxl-MUTANT        .......... .......... .......... .......... ..........
podxl-wildtype      TGATATCGTT TGTGATCTTA TTTTGTCTTT TGCAGCCTGT CTATGATCTG

podxl-MUTANT        .......... .......... .......... .......... ..........
podxl-wildtype      AGCAGCAGCA GCAGCAGCAC TGAAGTGAGT ACAGGAATGA AATTACAATG

podxl-MUTANT        .......... .......... .......... .......... ..........
podxl-wildtype      ACTGATGATG TCTTTAAACA TTCAGAGGTC ATAGAGATCA TTCATTTCTC

podxl-MUTANT        .......... .......... .......... .......... ..........
podxl-wildtype      TACCTGATGA CAATATGGCC TTTTTAAAAA GCATCATCCC CTTTAAAAAT

podxl-MUTANT        .......... .......... .......... .......... ..........
podxl-wildtype      AAAAAAAGTG AATAAATAAA TAAAAAAAAA CAGCATGTTC TCATCTGGAG

podxl-MUTANT        .......... .......... .......... .......... ..........
podxl-wildtype      GAAAAAAAAT CCATCTTATT TTCATCCTAA AATTCACCTG CACATATCAG

podxl-MUTANT        .......... .......... .......... .......... ..........
podxl-wildtype      GAATTTCTAT TATTTTTCAA TGTTCAAAGA CATCATTCAT TAATAAGAAA

podxl-MUTANT        .......... .......... .......... .......... ..........
podxl-wildtype      ATTTATTACA GAACAATGGA GCCAGAGGAT GTGTTTTTGT AAAACAATAG

podxl-MUTANT        .......... .......... .......... .......... ..........
podxl-wildtype      CATGCAAAGT AACTTAATTC ACAAAACTAT CTAACTTTCT CCTTAAAAAC
```

```
podxl-MUTANT        .......... .......... .......... .......... ..........
podxl-wildtype      ACTAAACACG CATTTGTGAT TGCAGAGGAA TATCGTGCGT GAAACCTGCA

podxl-MUTANT        .......... .......... .......... .......... ..........
podxl-wildtype      AGAAGCTGGG CCAGAACTTG AAAGGAAATT GCTCTGTGGA AGTGGAAATT

podxl-MUTANT        .......... .......... .......... .......... ..........
podxl-wildtype      AACAACAACC AATTAATTGC AACAATAACA ATAAATGGTG AGCAAAAGTT

podxl-MUTANT        .......... .......... .......... .......... ..........
podxl-wildtype      ACAGCTTCCA ATCATATAGA AGCACATCAT CTGGAGGAAT CACTTTTCCT

podxl-MUTANT        .......... .......... .......... .......... ..........
podxl-wildtype      CGTATTATCT CATCTTTAAT AACAATGAAA ATGATAGAAA TTCCTTGTTC

podxl-MUTANT        .......... .......... .......... .......... ..........
podxl-wildtype      ATTCGCAGCT AACCAGGCGA TACCATTGGA AAATTACAAT CCTCCAGAAA

podxl-MUTANT        .......... .......... .......... .......... ..........
podxl-wildtype      CAAATGTAAG CAGATTTAAC AGTAGCGCTT GCAATTTCCA AAGAAGTCTG

podxl-MUTANT        .......... .......... .......... .......... ..........
podxl-wildtype      TGCATTAAAA ACACCTTTTG TTTTCTTTGT AGGAAAACGA GAACAGCAAC

podxl-MUTANT        .......... .......... .......... .......... ..........
podxl-wildtype      AAGGAGCCAG TTCAAGACAC GATACCAGAC ACGCTGATTG CCATCTTGGC

podxl-MUTANT        .......... .......... .......... .......... ..........
podxl-wildtype      TTCATGTGGT GCTTTGGTGC TCATTCTGTG TGGCTTTGCT GCGTACTGCA

podxl-MUTANT        .......... .......... .......... ....AACACT CACCTCCTAA
podxl-wildtype      CTTATCATCG TAGATCCTAC AGGAAGAACC AGGTAACACT CACCTCCTAA

podxl-MUTANT        CACAAAGTTT ATAATGAAGG AACTCCAAGA ATGGCATCTA GATAGAGTAT
podxl-wildtype      CACAAAGTTT ATAATGAAGG AACTCCAAGA ATGGCATCTA GATAGAGTAT

podxl-MUTANT        AAAAACAGAT AGAATAATAA CAATGCAATT GTGTCCCAGA TCAGACACTG
podxl-wildtype      AAAAACAGAT AGAATAATAA CAATGCAATT GTGTCCCAGA TCAGACACTG

podxl-MUTANT        GGCATTCTGA AAGAGAAAGA CTTACTGTAA TCTTACCAAG ATAAATACTG
podxl-wildtype      GGCATTCTGA AAGAGAAAGA CTTACTGTAA TCTTACCAAG ATAAATACTG

podxl-MUTANT        GTTTTATGGA CCCGGAAAGG AACACACTTT GGCCTTTTAA AGTCTTAAAA
podxl-wildtype      GTTTTATGGA CCCGGAAAGG AACACACTTT GGCCTTTTAA AGTCTTAAAA

podxl-MUTANT        AACACCCTAA ACCAGGAGTG TTGAACTCTT CTGAGAATTT CTGGAGAGCC
```

```
podxl-wildtype       AACACCCTAA ACCAGGAGTG TTGAACTCTT CTGAGAATTT CTGGAGAGCC


podxl-MUTANT         ACAGCCCTGC ACACTTTAGT TCCAACCCAA CTCCAACAGA CCTGTAAGTT
podxl-wildtype       ACAGCCCTGC ACACTTTAGT TCCAACCCAA CTCCAACAGA CCTGTAAGTT


podxl-MUTANT         TCAAACAAGC CTGAAGGACT CAATTAGTTT GATCAGGTGT GTTTAATTAG
podxl-wildtype       TCAAACAAGC CTGAAGGACT CAATTAGTTT GATCAGGTGT GTTTAATTAG


podxl-MUTANT         GGTTGGAACT AAACTGCAGA GCTGCAACAA TGCAAGAACT GAGTTTGACA
podxl-wildtype       GGTTGGAACT AAACTGCAGA GCTGCAACAA TGCAAGAACT GAGTTTGACA


podxl-MUTANT         CCTGTGCCCT AAACCCTCAT GGTGCAAAGA TTGTATTTTT TTGCGAGTCA
podxl-wildtype       CCTGTGCCCT AAACCCTCAT GGTGCAAAGA TTGTATTTTT TTGCGAGTCA


podxl-MUTANT         AGCAGAACAG TTCTCTTAGC TTAACCTTTA CATTCCTTGT GGATCCAGGC
podxl-wildtype       AGCAGAACAG TTCTCTTAGC TTAACCTTTA CATTCCTTGT GGATCCAGGC


podxl-MUTANT         CAGTCAGCAC GGATACGGTT TCGTTTTCCA CTGGGGTGCT GATAACGGAG
podxl-wildtype       CAGTCAGCAC GGATACGGTT TCGTTTTCCA CTGGGGTGCT GATAACGGAG


podxl-MUTANT         CTCTCTGAAA TGTAATACAA ATGCGTCATG TGTTGCCTCG GTAATGCAAG
podxl-wildtype       CTCTCTGAAA TGTAATACAA ATGCGTCATG TGTTGCCTCG GTAATGCAAG


podxl-MUTANT         TGTAATGCAA ACACTGATAT GGATGGATAT CAAATATTTC TGTACAATAA
podxl-wildtype       TGTAATGCAA ACACTGATAT GGATGGATAT CAAATATTTC TGTACAATAA


podxl-MUTANT         GGTTGTGTTA GTTAATGCAT TTGCTAACAT GAGCAAATGA TGAACCACAC
podxl-wildtype       GGTTGTGTTA GTTAATGCAT TTGCTAACAT GAGCAAATGA TGAACCACAC


podxl-MUTANT         ATTTACTACA TTTAGTTAAT GAAAATGTTT GTTTGTTCAT GTTAACAAGT
podxl-wildtype       ATTTACTACA TTTAGTTAAT GAAAATGTTT GTTTGTTCAT GTTAACAAGT


podxl-MUTANT         GCATTAACTA ATGTTAAAAA GCATGAACTA GGATGTTAAC AATGCATTGC
podxl-wildtype       GCATTAACTA ATGTTAAAAA GCATGAACTA GGATGTTAAC AATGCATTGC


podxl-MUTANT         TAAATGTTGA ACTATGATAA ATAAACCCTA TACAAGTATT GTTCATAATT
podxl-wildtype       TAAATGTTGA ACTATGATAA ATAAACCCTA TACAAGTATT GTTCATAATT


podxl-MUTANT         AGTTCATGTT AGTAAATACA TGAACTAGTC AAATCTTAAT GTAAAGTGTG
podxl-wildtype       AGTTCATGTT AGTAAATACA TGAACTAGTC AAATCTTAAT GTAAAGTGTG


podxl-MUTANT         GCCATCATTC CCCCCCGTGT ATGCATATAT ACATGGATGT CAGAATCAGA
podxl-wildtype       GCCATCATTC CCCCCCGTGT ATGCATATAT ACATGGATGT CAGAATCAGA


podxl-MUTANT         ATTGGTTTTA TTGCCAAGTG TGCTTACGCA CACAAGAAAT TTGCTTTGGC
podxl-wildtype       ATTGGTTTTA TTGCCAAGTG TGCTTACGCA CACAAGAAAT TTGCTTTGGC
```

```
podxl-MUTANT         TACAGAAGCT TCCAGTGTGC ATAAAGTGAC AACACAAAAT AAATATGAGA
podxl-wildtype       TACAGAAGCT TCCAGTGTGC ATAAAGTGAC AACACAAAAT AAATATGAGA


podxl-MUTANT         AAACACGACA AATATTAAAC AGAGATGCAG TTAGACAATA TATAAGCATA
podxl-wildtype       AAACACGACA AATATTAAAC AGAGATGCAG TTAGACAATA TATAAGCATA


podxl-MUTANT         TTGTTATAAA TACACAAGTT ATATGGTGCT GTGTACAAAT GCATATGGAA
podxl-wildtype       TTGTTATAAA TACACAAGTT ATATGGTGCT GTGTACAAAT GCATATGGAA


podxl-MUTANT         AAAGTATTGC ATTGTACTGT ATATTATTAT AAAGTATGTG TGTATTTATA
podxl-wildtype       AAAGTATTGC ATTGTACTGT ATATTATTAT AAAGTATGTG TGTATTTATA


podxl-MUTANT         TAAAAAATAT GCACATTATT TATATGTACA TATTAAATAT ACACATTAAA
podxl-wildtype       TAAAAAATAT GCACATTATT TATATGTACA TATTAAATAT ACACATTAAA


podxl-MUTANT         AACTAAATTA CACAATTCAA ACTCTTATTT ATGTGATTGG TCGTGATTAA
podxl-wildtype       AACTAAATTA CACAATTCAA ACTCTTATTT ATGTGATTGG TCGTGATTAA


podxl-MUTANT         CAGCACGATT AACATCCCAA TGTGTACTTT TTACCTGGAC AGCAACATCT
podxl-wildtype       CAGCACGATT AACATCCCAA TGTGTACTTT TTACCTGGAC AGCAACATCT


podxl-MUTANT         AACAGAAGAG CTGCAGACTG TGGAAAACGG TTATCATGAC AATCCCACTC
podxl-wildtype       AACAGAAGAG CTGCAGACTG TGGAAAACGG TTATCATGAC AATCCCACTC


podxl-MUTANT         TGGAGGTGAT GGAGGTGCAG CCTGAGATGC AAGAGAAGAA ACTGGCATTG
podxl-wildtype       TGGAGGTGAT GGAGGTGCAG CCTGAGATGC AAGAGAAGAA ACTGGCATTG


podxl-MUTANT         AATGGAGAGT TTAACGACAG CTGGATCGTC CCAATAGACA ACCTTCTGAA
podxl-wildtype       AATGGAGAGT TTAACGACAG CTGGATCGTC CCAATAGACA ACCTTCTGAA


podxl-MUTANT         AGAGGACATA CCTGACGAAG AGGACACTCA CTTGTAATGG ACCCCGGTGA
podxl-wildtype       AGAGGACATA CCTGACGAAG AGGACACTCA CTTGTAATGG ACCCCGGTGA


podxl-MUTANT         GCCGTGACGA GTGGGAAACC TGCTGCTGCC TTTACGGTAA ATTGTGACCA
podxl-wildtype       GCCGTGACGA GTGGGAAACC TGCTGCTGCC TTTACGGTAA ATTGTGACCA


podxl-MUTANT         GCCAGAATGT CAAGAAAAGC AGAGATCGAC GTAAAGACTA TGACAAGATA
podxl-wildtype       GCCAGAATGT CAAGAAAAGC AGAGATCGAC GTAAAGACTA TGACAAGATA


podxl-MUTANT         GTGGTTTCAT TTGGCGCCCA CACTGTATAT GCTGTAATTA AATAGTTTGA
podxl-wildtype       GTGGTTTCAT TTGGCGCCCA CACTGTATAT GCTGTAATTA AATAGTTTGA


podxl-MUTANT         TGTCAAGTAT GTGCTAGAAA GAGTAGAAAA GCTTTTGAGT TTTGTGTGTC
podxl-wildtype       TGTCAAGTAT GTGCTAGAAA GAGTAGAAAA GCTTTTGAGT TTTGTGTGTC


podxl-MUTANT         ACTGTCTTTG GAAATGATCA ATGCACAATC TATGGCCTCT AATTGTGCCT
podxl-wildtype       ACTGTCTTTG GAAATGATCA ATGCACAATC TATGGCCTCT AATTGTGCCT
```

```
podxl-MUTANT        TACTTTTCTT CATCACACGG TGGAGAAACC AGTAGTTTGA TACGCATACA
podxl-wildtype      TACTTTTCTT CATCACACGG TGGAGAAACC AGTAGTTTGA TACGCATACA

podxl-MUTANT        AATATATACA CAGACGGATG TTGGGGATCA GCCTGTTTTA TTATTTTGAA
podxl-wildtype      AATATATACA CAGACGGATG TTGGGGATCA GCCTGTTTTA TTATTTTGAA

podxl-MUTANT        CACTTTTGGT TCAGTTATTA TTAATATTGA ATGAAAAAAA ACAGGATTAT
podxl-wildtype      CACTTTTGGT TCAGTTATTA TTAATATTGA ATGAAAAAAA ACAGGATTAT

podxl-MUTANT        AATTTTATTG TAATTTCACT CACTATATTT GAACATTTTT TGCAACACAT
podxl-wildtype      AATTTTATTG TAATTTCACT CACTATATTT GAACATTTTT TGCAACACAT

podxl-MUTANT        TTTGAGACTT GTGAAGAAAT GATGCCATTT TTATCTAAAA ACAAGGAATC
podxl-wildtype      TTTGAGACTT GTGAAGAAAT GATGCCATTT TTATCTAAAA ACAAGGAATC

podxl-MUTANT        CGTTTTATTA TTTTTAACAA AACAAGTGTT TTTAAAGTGA AAGTTTGTGT
podxl-wildtype      CGTTTTATTA TTTTTAACAA AACAAGTGTT TTTAAAGTGA AAGTTTGTGT

podxl-MUTANT        CTGAAACAAA GAAACGTGGC TTTGTGTAAT TGGTGACGTA AAATAGAAGG
podxl-wildtype      CTGAAACAAA GAAACGTGGC TTTGTGTAAT TGGTGACGTA AAATAGAAGG

podxl-MUTANT        ACGACAGGAC CAAAACATTC AATGCACATT AACTACAGGA TCATATTTTA
podxl-wildtype      ACGACAGGAC CAAAACATTC AATGCACATT AACTACAGGA TCATATTTTA

podxl-MUTANT        ACTAGTGATG TTGCCACCTA CTGGCCTGGC ATGCATAATA CAGCCTTTTT
podxl-wildtype      ACTAGTGATG TTGCCACCTA CTGGCCTGGC ATGCATAATA CAGCCTTTTT

podxl-MUTANT        TTGATACAGT GGTCCCTCAT TATTCACGGG AGTGGAGTTC TAAAAATAAC
podxl-wildtype      TTGATACAGT GGTCCCTCAT TATTCACGGG AGTGGAGTTC TAAAAATAAC

podxl-MUTANT        CCGCTATTGG TAAAATCCGC AAAATAGTCT GCTTTATCTT TATCAATAAG
podxl-wildtype      CCGCTATTGG TAAAATCCGC AAAATAGTCT GCTTTATCTT TATCAATAAG

podxl-MUTANT        TATAGATGGT TTAAGGCTGT AAAACCCCTC ACTACACTAT TTTTCCGACA
podxl-wildtype      TATAGATGGT TTAAGGCTGT AAAACCCCTC ACTACACTAT TTTTCCGACA

podxl-MUTANT        GGCATTTTCA GACTTTTCTC TCGTTTAAAC TTCCATCCTC TTTTAGCATG
podxl-wildtype      GGCATTTTCA GACTTTTCTC TCGTTTAAAC TTCCATCCTC TTTTAGCATG

podxl-MUTANT        TAGAGAATTC CAACTTTTTG TGCGATGGTT AGCATCTTCC TTTGCTTTTT
podxl-wildtype      TAGAGAATTC CAACTTTTTG TGCGATGGTT AGCATCTTCC TTTGCTTTTT

podxl-MUTANT        GGGTGCTACC ATGGGTGCCT TTGATTATGC AGAATGTTTC GTCAGCAATA
podxl-wildtype      GGGTGCTACC ATGGGTGCCT TTGATTATGC AGAATGTTTC GTCAGCAATA

podxl-MUTANT        TGGGGATTGT TGGGGAGAAA ACTTGCAACA TACAGAACAG CACTTTAGAG
podxl-wildtype      TGGGGATTGT TGGGGAGAAA ACTTGCAACA TACAGAACAG CACTTTAGAG
```

```
podxl-MUTANT         TCACACTGCT AGCGATCAAA ATTTGGCAAG CTGAACGCAT TCTGTACTGT
podxl-wildtype       TCACACTGCT AGCGATCAAA ATTTGGCAAG CTGAACGCAT TCTGTACTGT

podxl-MUTANT         ACATGTGACA CGGCAAGGAG GAGATTGACT GGCAATGGTC TACAGCCAAT
podxl-wildtype       ACATGTGACA CGGCAAGGAG GAGATTGACT GGCAATGGTC TACAGCCAAT

podxl-MUTANT         CAGGACACAG AACACATTGT ACTAAACAGT AAGCAGAGAA CAATGCGCTG
podxl-wildtype       CAGGACACAG AACACATTGT ACTAAACAGT AAGCAGAGAA CAATGCGCTG

podxl-MUTANT         TAAAAAAAAA AAAAAAAAGC ATGTCAAATT ACACAAAAAA AACAGCGAAA
podxl-wildtype       TAAAAAAAAA AAAAAAAAGC ATGTCAAATT ACACAAAAAA AACAGCGAAA

podxl-MUTANT         CAGCAGACCG CAAAAGGTGA ACCGTGTTAT GGCGAGGGAC CACTGTAGTT
podxl-wildtype       CAGCAGACCG CAAAAGGTGA ACCGTGTTAT GGCGAGGGAC CACTGTAGTT

podxl-MUTANT         GGAGGATCTT TGTAAATAGG GATTTTTTAA AACCTTTTTA AAATGTACAG
podxl-wildtype       GGAGGATCTT TGTAAATAGG GATTTTTTAA AACCTTTTTA AAATGTACAG

podxl-MUTANT         AAACAAATTA CCAGTTTTAT GCACAATATT GTCATCTAAA ACAGTGTTTC
podxl-wildtype       AAACAAATTA CCAGTTTTAT GCACAATATT GTCATCTAAA ACAGTGTTTC

podxl-MUTANT         TAAACCACGT TCCTGGAGGA CCACCAGCAC TGCATGTTTT GGACGTCTCC
podxl-wildtype       TAAACCACGT TCCTGGAGGA CCACCAGCAC TGCATGTTTT GGACGTCTCC

podxl-MUTANT         GATGTCTGTC ACACTGATTA CAGCTTTCAG TCTCTGATTA TGAGCTGTTG
podxl-wildtype       GATGTCTGTC ACACTGATTA CAGCTTTCAG TCTCTGATTA TGAGCTGTTG

podxl-MUTANT         ATCTGAATCA GGTGTGTTTG GTTAAGGAGA CATGGATAAT GTGCAGAGCT
podxl-wildtype       ATCTGAATCA GGTGTGTTTG GTTAAGGAGA CATGGATAAT GTGCAGAGCT

podxl-MUTANT         GGTGGTCCCA CAGGATTGTG GTTGAGAAAC ACTGATCTAA ACACTCTAAA
podxl-wildtype       GGTGGTCCCA CAGGATTGTG GTTGAGAAAC ACTGATCTAA ACACTCTAAA

podxl-MUTANT         TCTGATGCAT TTGTTCTAAA GAAGGCCTCT TGGGAGACAT TATGCATTAA
podxl-wildtype       TCTGATGCAT TTGTTCTAAA GAAGGCCTCT TGGGAGACAT TATGCATTAA

podxl-MUTANT         AACAGTGTAA AGCACCATCA GGTTCTATTT TTACATGCAC ATTGCTCTCA
podxl-wildtype       AACAGTGTAA AGCACCATCA GGTTCTATTT TTACATGCAC ATTGCTCTCA

podxl-MUTANT         GTATTCCTCT TAAGTGTTTT AGCTTGCGGA TCACTGGATT CCCATTAAAT
podxl-wildtype       GTATTCCTCT TAAGTGTTTT AGCTTGCGGA TCACTGGATT CCCATTAAAT

podxl-MUTANT         TATGCTATAG TGGTTGTATC TACCAGAGAA CGAAACACAG TTTATAAATG
podxl-wildtype       TATGCTATAG TGGTTGTATC TACCAGAGAA CGAAACACAG TTTATAAATG

podxl-MUTANT         TTTTGTTATT AGAAACCTTT CCAGCCATAT TTTTAAGACT TGGACATTAT
```

```
podxl-wildtype       TTTTGTTATT AGAAACCTTT CCAGCCATAT TTTTAAGACT TGGACATTAT


podxl-MUTANT         ATTAATCCAG GACAGTTGGT ACCCTGCTTT ATTTCTTGAC AGGACTGTAA
podxl-wildtype       ATTAATCCAG GACAGTTGGT ACCCTGCTTT ATTTCTTGAC AGGACTGTAA


podxl-MUTANT         CTGCTCTTGT ATATAAGAAT TATTTTCTAT GCGTCTTCCA TGTTTTAGCC
podxl-wildtype       CTGCTCTTGT ATATAAGAAT TATTTTCTAT GCGTCTTCCA TGTTTTAGCC


podxl-MUTANT         CTCACTCACA GGCACCAAAT TCCACTGAGA CCAATGGATA ACTCATCCTG
podxl-wildtype       CTCACTCACA GGCACCAAAT TCCACTGAGA CCAATGGATA ACTCATCCTG


podxl-MUTANT         GTAAAATTAG ACGATAGTTT GAGAGATTGT GGCTTTCTGA TTGGCAGATA
podxl-wildtype       GTAAAATTAG ACGATAGTTT GAGAGATTGT GGCTTTCTGA TTGGCAGATA


podxl-MUTANT         ACCTTTTGCA TTATCTTCCC TCAGCGCTAC ATCTCTGTGC TGGTAGTCAT
podxl-wildtype       ACCTTTTGCA TTATCTTCCC TCAGCGCTAC ATCTCTGTGC TGGTAGTCAT


podxl-MUTANT         GCTGGTATGA AAATTTACCA TCTTGTCAAC CTGAAACATT TGACACTAAG
podxl-wildtype       GCTGGTATGA AAATTTACCA TCTTGTCAAC CTGAAACATT TGACACTAAG


podxl-MUTANT         TCCAACACAC ACATTGTGTA GCTTTTCATG TATTAAATCG TGCTGTACTG
podxl-wildtype       TCCAACACAC ACATTGTGTA GCTTTTCATG TATTAAATCG TGCTGTACTG


podxl-MUTANT         CCATTTGAAC TGGATTTTTG GTTTAAGTAC GCTGATCGCC TTTTTCGTTG
podxl-wildtype       CCATTTGAAC TGGATTTTTG GTTTAAGTAC GCTGATCGCC TTTTTCGTTG


podxl-MUTANT         TATTTAAGTG TTCTAAAGTG GTTTTCACCA AAACATTAAA TGTTTTCTAT
podxl-wildtype       TATTTAAGTG TTCTAAAGTG GTTTTCACCA AAACATTAAA TGTTTTCTAT


podxl-MUTANT         TATTATTTTT GTTGTATTTG CATGCACTAC ACAAATTGTC AATATAAGGG
podxl-wildtype       TATTATTTTT GTTGTATTTG CATGCACTAC ACAAATTGTC AATATAAGGG


podxl-MUTANT         ATTCACTGCT GCCTCCTGAT CTACTTCCAC CAAAGTATTA AGGTTGAATT
podxl-wildtype       ATTCACTGCT GCCTCCTGAT CTACTTCCAC CAAAGTATTA AGGTTGAATT


podxl-MUTANT         GATTGCTGTA TTCTGTAATT GAAGTCAGTT TGCTACTGTG TGCAAATGAC
podxl-wildtype       GATTGCTGTA TTCTGTAATT GAAGTCAGTT TGCTACTGTG TGCAAATGAC


podxl-MUTANT         AGGGGTCTGA TACACTGGAA GCCGCATCTC TGTCAGATCA GACGGGGACG
podxl-wildtype       AGGGGTCTGA TACACTGGAA GCCGCATCTC TGTCAGATCA GACGGGGACG


podxl-MUTANT         GCCATGATTT CTAATACAAA ATAAAAAATG TGATTTTTTT TTTAAA
podxl-wildtype       GCCATGATTT CTAATACAAA ATAAAAAATG TGATTTTTTT TTTAAA
```

**S6 Fig. *podxl*$^{Ex1(p)\_Ex7\Delta}$ mutant DNA sequence.** Expected *podxl*$^{Ex1(p)_Ex7\Delta}$mutant DNA sequence was aligned to wildtype using T-coffee (Notredame et al. 2000).

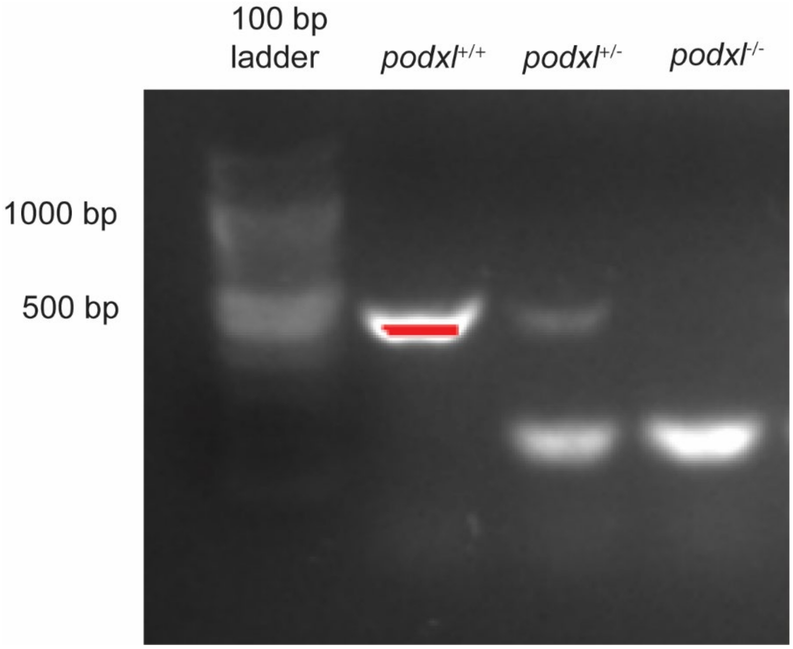


**S7 Fig. *podxl*$^{Ex1(p)\_Ex7\Delta}$ mutant PCR.** To genotype *podxl*$^{Ex1(p)\_Ex7\Delta}$ mutants, PCR was performed on wildtype, heterozygous mutant, and homozygous mutant fish using primers from S8 Table. Primers are expected to amplify a 450 bp fragment from the wildtype allele (lanes 2 and 3) and a 155 bp fragment from the mutant allele (lanes 3 and 4).

```
podxl-Transmembrane-Actual      GTGAGTCTTT CTCTTTCAGA ATGCCCAGTG TCTGATCTGG GACACAATTG
podxl-Transmembrane-Expected    GTAAGTCTTT CTCTTTCAGA ATGCCCAGTG TCTGATCTGG GACACAATTG


podxl-Transmembrane-Actual      CATTGTTATT ATTCTATCTG TTTTTATACT CTATCTAGAT GCCATTCTTG
podxl-Transmembrane-Expected    CATTGTTATT ATTCTATCTG TTTTTATACT CTATCTAGAT GCCATTCTTG


podxl-Transmembrane-Actual      GAGTTCCTTC ATTATAAACT TTGTGTTAGG AGGTGAGTGT TCAATCACAA
podxl-Transmembrane-Expected    GAGTTCCTTC ATTATAAACT TTGTGTTAGG AGGTGAGTGT TCAATCACAA


podxl-Transmembrane-Actual      ATGCGTGTTT ACTGTTTTTA AGGAGA
podxl-Transmembrane-Expected    ATGCGTGTTT AGTGTTTTTA AGGAGA
```

**S8 Fig. *podxl*$^{Ex5\_Ex7\Delta}$ mutant RNA sequence.** *podxl* mutant RNA sequence from Sanger sequencing was aligned to expected sequence using T-coffee (Notredame et al. 2000). The pink highlighted region is the cut site for the two sgRNAs.

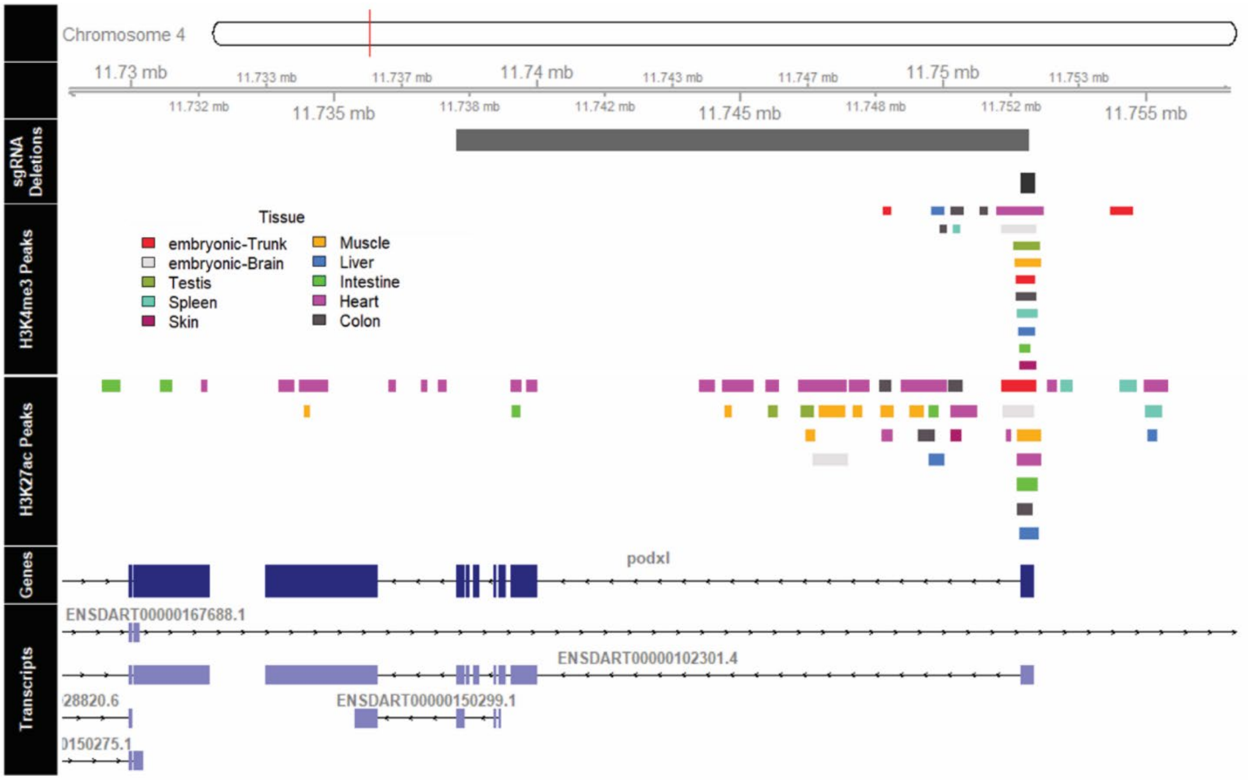


**S9 Fig. Histone modifications near *podxl* throughout zebrafish tissues and development.** The region of zebrafish chromosome 4 containing *podxl* +/- 5kb is shown. Both promoter deletion alleles ($podxl^{-194\_Ex7\Delta}$ and $podxl^{-319\_Ex1(p)\Delta}$) allele are named according to the approximate cut sites of the CRISPR targeting sgRNAs relative to the *podxl* coding sequence. The $podxl^{-194\_Ex7\Delta}$ allele was generated by targeting chr4:11,738,026 and chr4:11,752,120 and is shown in grey. The $podxl^{-319\_Ex1(p)\Delta}$ allele was generated by targeting chr4:11,751,916 and chr4:11,752,245 and is shown in black. H3K4me3 peaks in this region (Yang et al. 2020), characteristic of promoters, are shown in the panel below the deletion sites and colored by tissue and developmental stage. Similarly, H3K27ac peaks (Yang et al. 2020), are shown and colored identically. The position of these features is plotted above the *podxl* refGene model (dark blue) and Ensembl transcripts (light blue) within this region. Both deletion alleles overlap with the canonical transcriptional start site identified in every tissue with H3K4me3 peaks along this gene. The $podxl^{-194\_Ex7\Delta}$ allele additionally removes H3K4me3 peaks detected within intron 1 of *podxl*. However, several likely enhancer peaks (indicated by H3K27ac marks without H3K4me3) remain in both alleles and a single upstream promoter peak (indicated by H3K4me3) was detected in tissue from the embryonic trunk. It is unclear if this retained H3K4me3 peak is associated with *podxl* transcription or other nearby genetic elements.

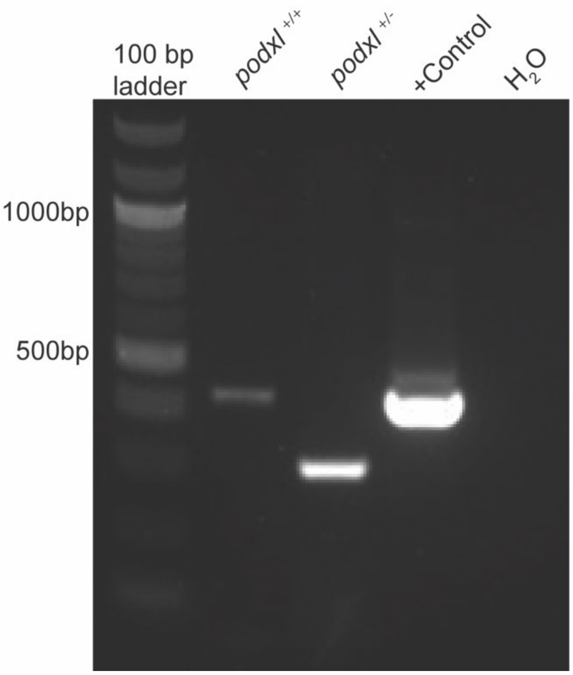


**S10 Fig. *podxl*$^{-194\_ex7\Delta}$ mutant PCR.** To genotype *podxl*$^{-194\_ex7\Delta}$ mutants, PCR was performed on wildtype and heterozygous mutant fish using primers from S8 Table. Primers are expected to amplify a 492 bp fragment from the wildtype allele (lanes 2 and 4) and a 292 bp fragment from the mutant allele (lane 3).

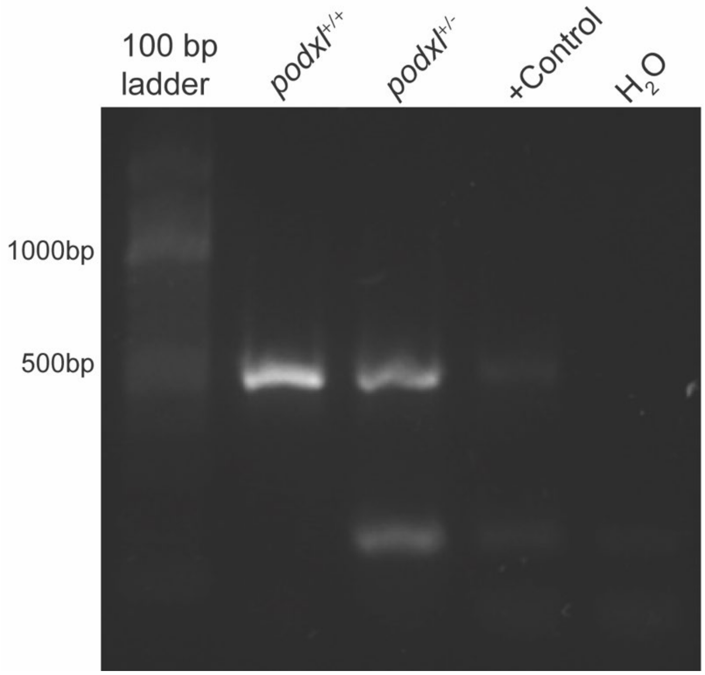


**S11 Fig.** ***podxl$^{-319_ex1(p)\Delta}$*** **mutant PCR.** To genotype *podxl$^{-319_ex1(p)\Delta}$* mutants, PCR was performed on wildtype and heterozygous mutant fish using primers from S8 Table. Primers are expected to amplify a 492 bp fragment from the wildtype allele (lanes 2-4) and a 163 bp fragment from the mutant allele (lanes 3 and 4).

```
Podxl-Mutant          GCAAACAAAA GCTCTCCTCT TGGCTTTCAG ATAAAAACAT ACCAGCTGTG
Podxl-Wildtype        GCAAACAAAA GCTCTCCTCT TGGCTTTCAG ATAAAAACAT ACCAGCTGTG


Podxl-Mutant          AAAGAAAAAA AAAGTCCTGA TACAAACTCA AGACGAAAAG CGGAACCGAG
Podxl-Wildtype        AAAGAAAAAA AAAGTCCTGA TACAAACTCA AGACGAAAAG CGGAACCGAG


Podxl-Mutant          CGGCGGATCC ACGTTTCCCA TGCTAAACTG TTGCTGAAGA GACTGAACGC
Podxl-Wildtype        CGGCGGATCC ACGTTTCCCA TGCTAAACTG TTGCTGAAGA GACTGAACGC


Podxl-Mutant          GGAGAATCTG AAAATGACCA TCA....... .......... ..........
Podxl-Wildtype        GGAGAATCTG AAAATGACCA TCACGTGGAC AATCATCGTA TTAGGTGAGT


Podxl-Mutant          .......... .......... .......... .......... ..........
Podxl-Wildtype        TTACCTTCAT ATACGACGAG GATGCAGCTT AAATATAAAC TCGATCTCTT


Podxl-Mutant          .......... .......... .......... .......... ..........
Podxl-Wildtype        CAGTAATTAA ATATAATGTT AGTCGGTTAA TTTGAGGGAC ATGTGCTGGG
```

**S12 Fig. *podxl* sgRNA #1 cut site is absent in *podxl*$^{Ex1(p)\_Ex7\Delta}$ mutants.** We used T-coffee (Notredame et al. 2000) to align sequences in *podxl* mutant and wildtype. The cut site is highlighted in pink and the rest of the sgRNA site is highlighted in blue.

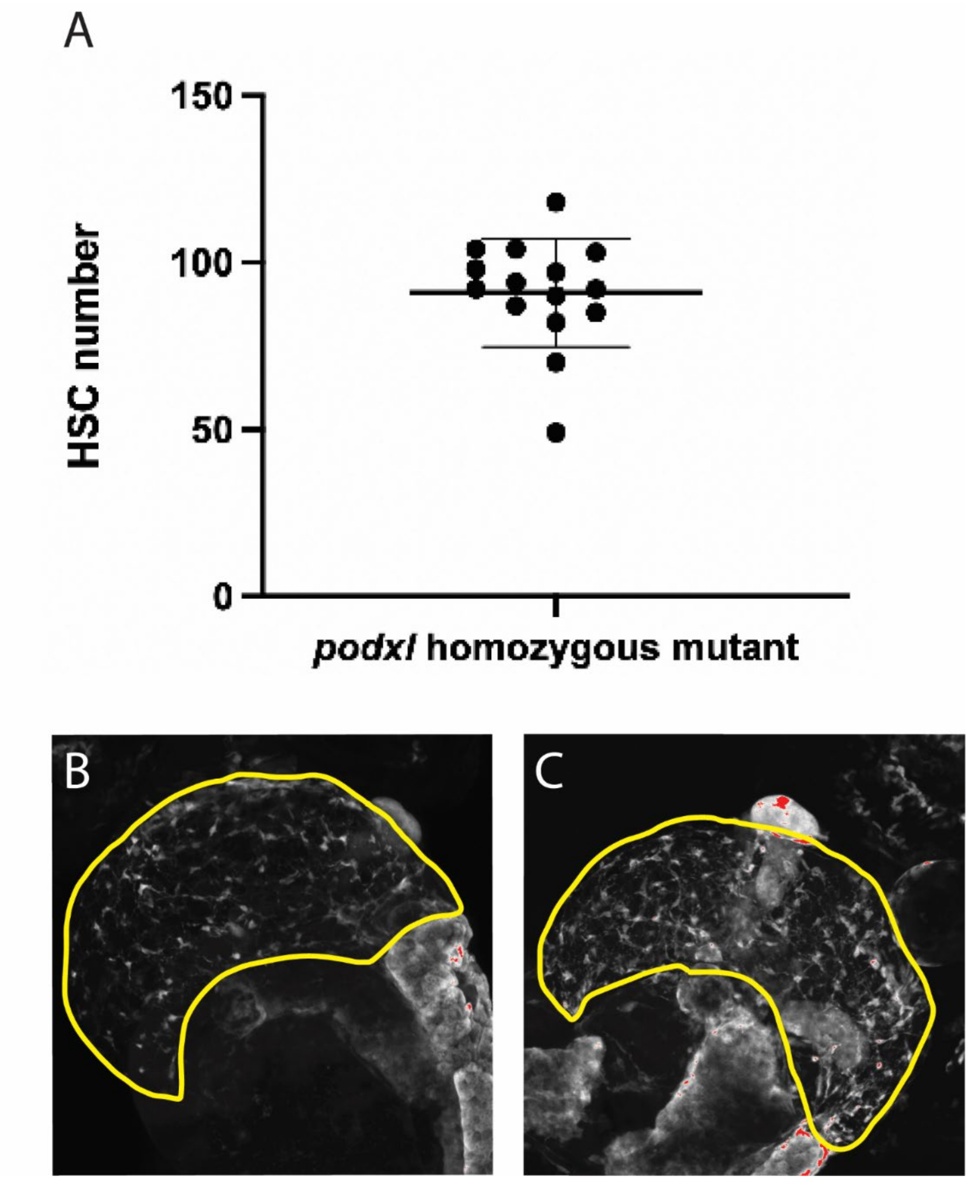


**S13 Fig. Maternal zygotic *podxl* mutants show unremarkable HSC morphology and number.** Progeny from a *podxl*$^{Ex1(p)\_Ex7\Delta}$ homozygous mutant zebrafish incross in the *Tg(wt1b):EGFP* background were fixed at 6 dpf, stained with anti-EGFP antibodies, imaged with confocal microscopy, and analyzed by manual examination and counting of HSCs. HSC numbers (A) and morphology (B and C) were similar to those observed in wildtype zebrafish (Fig 1 and 2). Bars show mean +/- S.D.

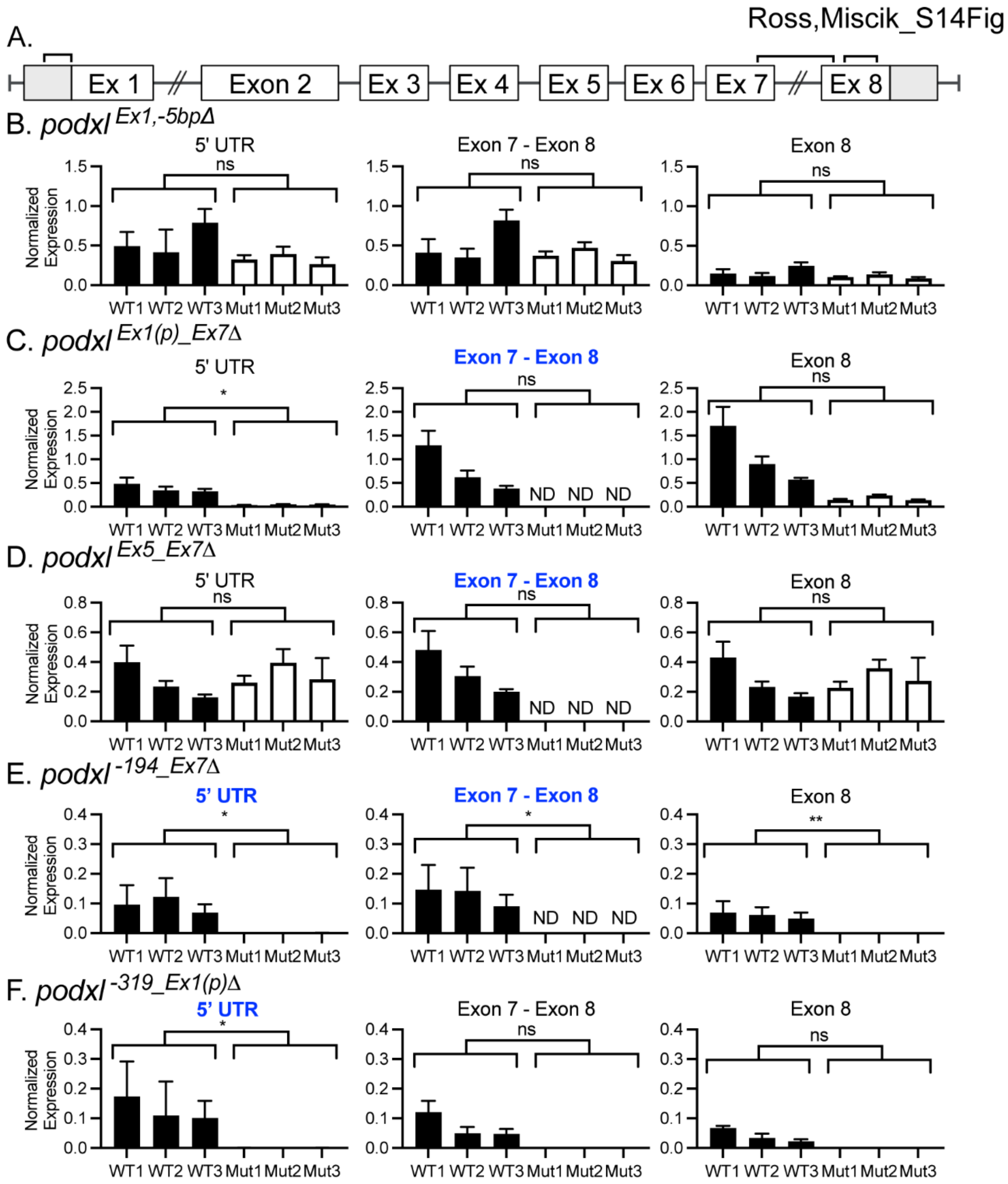


**S14 Fig. Expression of *podxl* mRNA in *podxl* mutants.** These are the same data as in Figure 4, but presented differently to show technical variation. Here each bar represents the mean +/- SD of 2-3 technical replicates (distinct qPCR reactions). (A) Wildtype *podxl* gene schematic with brackets indicating the regions of mRNA amplified by qPCR; 5'UTR - exon 1(left), exon 7 - exon 8 (center), and partial exon 8 (right). Normalized expression levels of each region from the livers of 3mpf *podxl* $^{Ex1,-5bp\Delta}$ (B), *podxl*$^{\mathrm{Ex1(p)\_Ex7}\Delta}$ (C), *podxl*$^{\mathrm{Ex5\_Ex7}\Delta}$ (D), *podxl*$^{\mathrm{-194\_Ex7}\Delta}$ (E), *podxl*$^{\mathrm{-319\_Ex1(p)}\Delta}$ (F) mutants and wildtype siblings. Primers lacking binding sites in the mutant allele are indicated by bold blue graph titles. Welch's t-test. NS, not significant; *, p<0.05; **, p<0.01; ND, not detected.

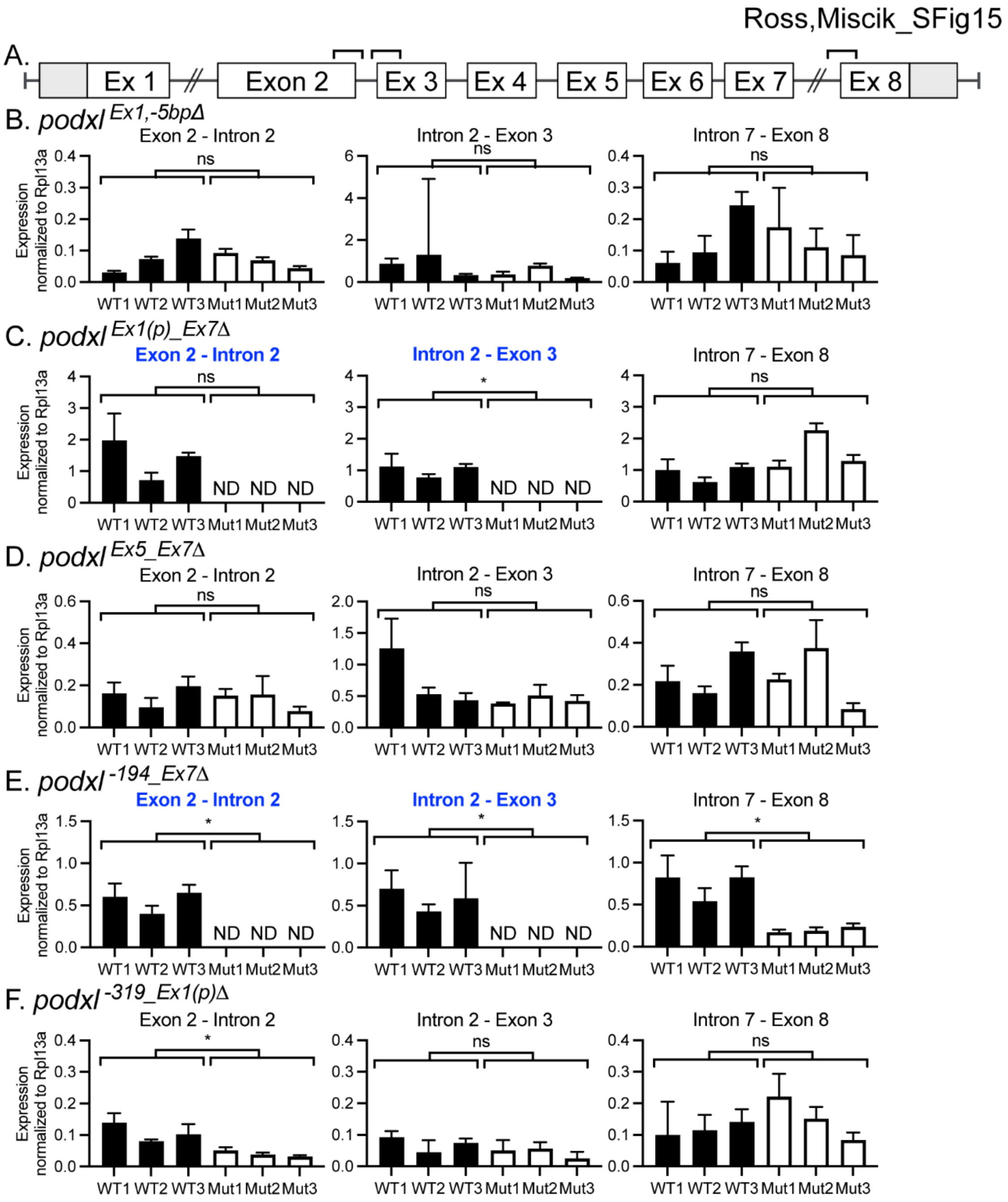


**S15 Fig. Expression of *podxl* pre-mRNA in *podxl* mutants.** These are the same data as in Figure 5, but presented differently to show technical variation. Here each bar represents the mean +/- SD of 2-3 technical replicates (distinct qPCR reactions). (A) Wildtype *podxl* gene schematic with brackets indicating the regions of pre-mRNA amplified by qPCR; exon 2 - intron 2 (left), intron 2 - exon 3 (center), and intron 7 - exon 8 (right). ). Normalized expression levels of each region from the livers of 3mpf $podxl^{Ex1,-5bp\Delta}$ (B), $podxl^{Ex1(p)\_Ex7\Delta}$ (C), $podxl^{Ex5\_Ex7\Delta}$ (D), $podxl^{-194\_Ex7\Delta}$ (E), $podxl^{-319\_Ex1(p)\Delta}$ (F) mutants and wildtype siblings. Primers lacking binding sites in the mutant allele are indicated by bold blue graph titles. Welch’s t test. NS, not significant; *, p<0.05.

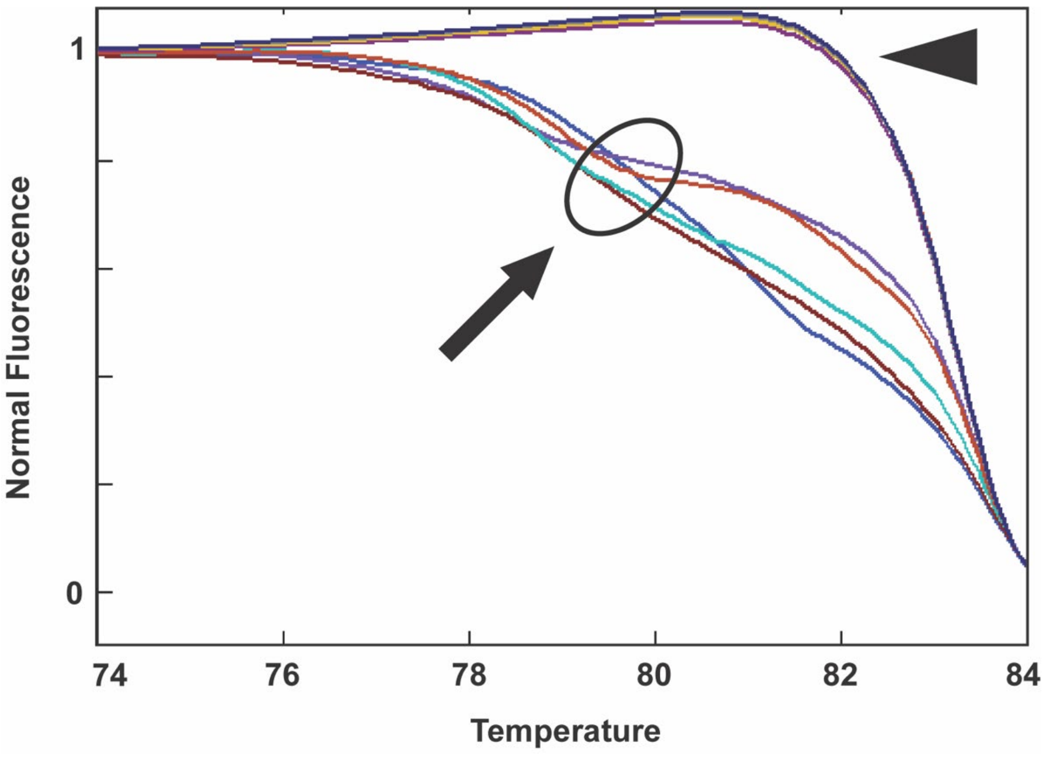


**S16 Fig. *endo* HRMA confirmation.** The presence of *endo* mutations in every injected embryo was confirmed by amplifying the region surrounding the CRISPR target site and performing high-resolution melt analysis (HRMA). Wild-type *endo* (arrowhead) and mutated *endo* (circled, arrow) show distinct shifted melting curves. Only zebrafish with confirmed mutations were included in data analysis.

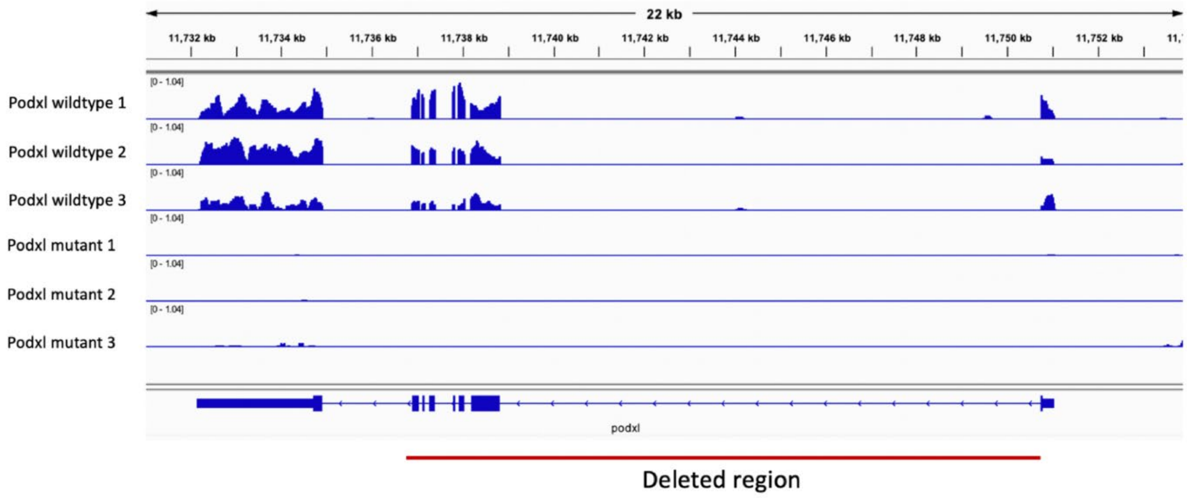


**S17 Fig. RNA sequencing reads for *podxl* gene in wildtype and *podxl*$^{Ex1(p)\_Ex7\Delta}$ mutants.** The red bar shows region that is deleted in *podxl*$^{Ex1(p)\_Ex7\Delta}$ mutants.

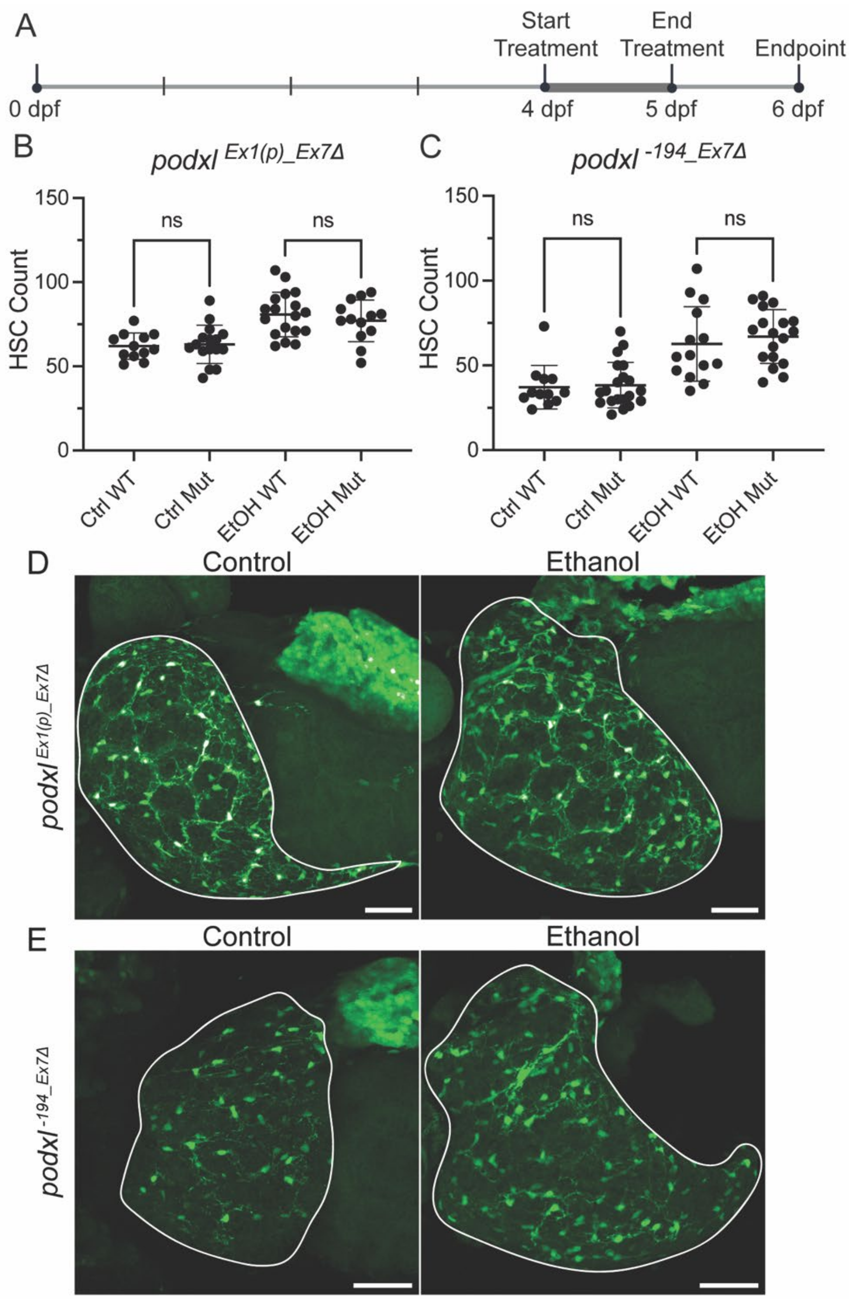


**S18 Fig. *podxl* mutants respond similarly to wildtype zebrafish in response to ethanol treatment.** *Podxl* mutants were exposed to 2% ethanol in egg water or egg water alone following the timeline (A). The HSC count was evaluated for *podxl* $^{Ex1(p)\_Ex7\Delta}$ (B) and *podxl* $^{-194\_Ex7\Delta}$ (C) and wildtype control siblings. (D,E) Representative images; Scale bars are 50 µm. Bars show mean +/- SD. Ordinary one-way ANOVA (B) and Kruskal-Wallis test (C). NS, not significant.

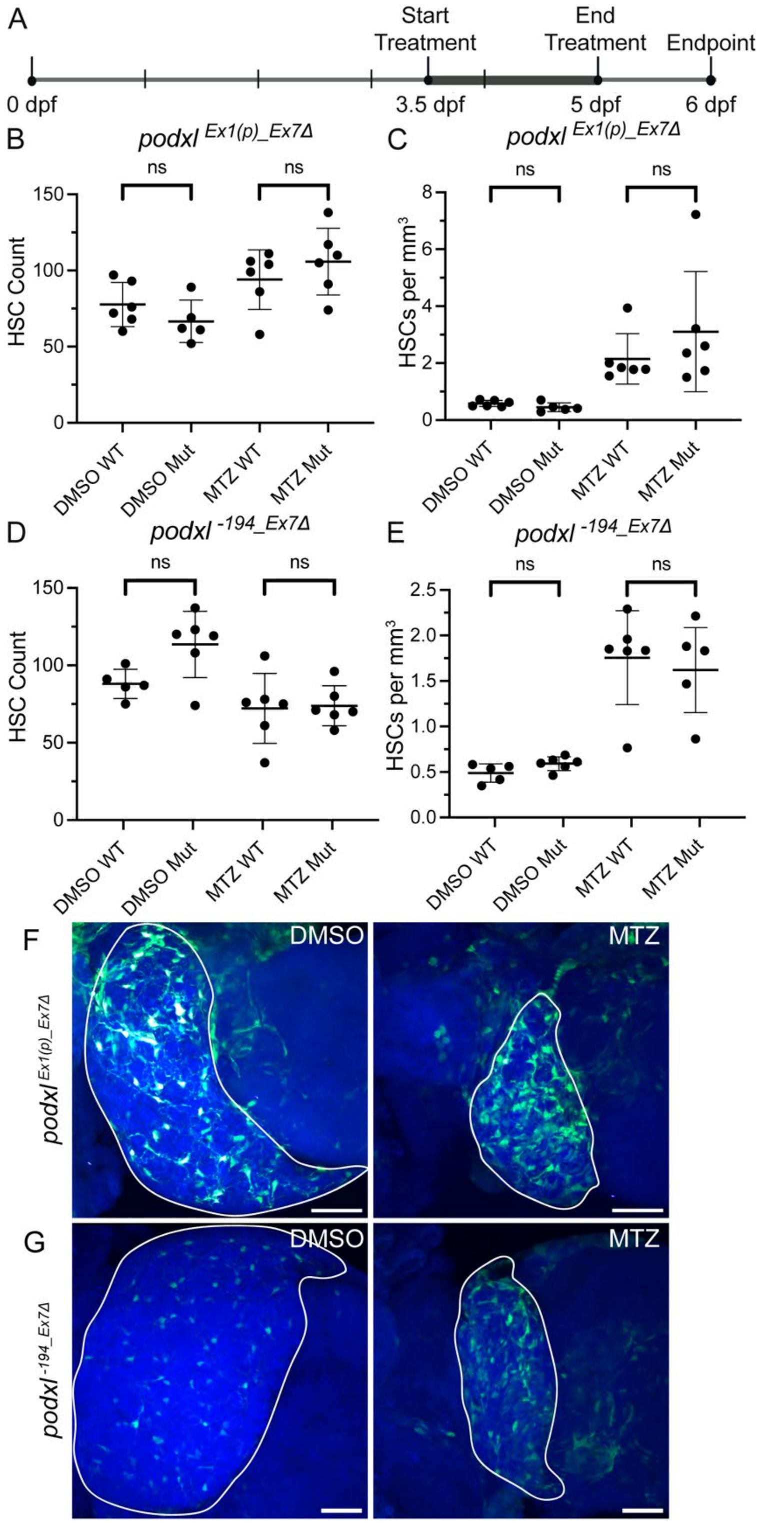


803 **S19 Fig. *podxl* mutants respond similarly to wildtype zebrafish in response to hepatocyte ablation.** *Podxl* mutants were exposed to MTZ or DMSO following the timeline (A). The HSC count and HSC density were evaluated for *podxl* $^{Ex1(p)\_Ex7\Delta}$ (B,C) and *podxl* $^{-194\_Ex7\Delta}$ (D,E) and

wildtype control siblings. (F,G) Representative images; Scale bars are 50 μm. Bars show mean +/- SD. Ordinary one-way ANOVA (B,D) and Kruskal-Wallis test (C,E). NS, not significant.

### Supplementary Tables

**S1 Table. Sequences used for *podxl, endo,* and *tyr* knockdown and *podxl* morpholino.**

| **Single guides for CRISPR** | **Strand** | **Sequence** |
|---|---|---|
| *podxl* CRISPR 1-exon 1 | + | TACGATGATTGTCCACGTGATGG |
| *podxl* CRISPR 2-exon 2 | + | AGTGACCACACTTCTATTCGAGG |
| *tyr* CRISPR-exon1 | - | GGACTGGAGGACTTCTGGGGTGT |
| *endo* CRISPR-exon 1 | - | AGAGTTCCCAGGAGAGCTCCGGG |
| **Morpholino**<br>*podxl* | | GGT CAT TTT CAG ATT CTC CGC GTT C |

**S2 Table. qPCR primers**

| RT-PCR primers | Used for | | |
|---|---|---|---|
| **Gene-exon** | | **Forward Sequence** | **Reverse Sequence** |
| B-actin-exon 3-4 | Normalization | CTATGAGCTGCCTGACGGTCA | GTGGTCTCGTGGATACCGCAA |
| Rpl13a | Normalization | TAAGGACGGAGTGAACAACCA | CTTACGTCTGCGGATCTTTCTG |
| Podxl-5'UTR | mRNA | ACACACCAAAACTTTGCCCC | TCGGTTCCGCTTTTCGTCTT |
| Podxl-exon7-8 | mRNA | CATGATAACCGTTTTCCACAGTCTG | GCTCATTCTGTGTGGCTTTGC |
| Podxl-exon8<br>(exon 8-3'UTR) | mRNA | GCAGCCTGAGATGCAAGAGA | TTACCGTAAAGGCAGCAGCA |
| Podxl-exon2-intron2 | Pre-mRNA | CAGCCACACCAGCTAACTCT | TCATTCTGATGTGAGCGAAATCTT |
| Podxl-intron2-exon3 | Pre-mRNA | AGATTTCGCTCACATCAGAATGA | TTTGGTTTTGCTGTGGTGAGTA |
| Podxl-intron7-exon8 | Pre-mRNA | AACAGCACGATTAACATCCCAA | GGACGATCCAGCTGTCGTTA |

**S3 Table. Top 100 upregulated genes and genes of interest in *podxl*$^{Ex1(p)\_Ex7\Delta}$ mutants found using RNA sequencing.** Extracellular region genes are highlighted in yellow, extracellular matrix genes are highlighted in blue, and genes that are in both categories are highlighted in green.

| Number based on Log2 fold change | Gene name | Log2 fold change | Adjusted p value |
|---|---|---|---|
| 1 | *cyp2k22* | 10.8596734 | 1.0299E-46 |
| 2 | *hsd17b3* | 8.39101201 | 1.4462E-44 |
| 3 | *cd59* | 8.33176546 | 4.6412E-15 |
| 4 | *slc27a6* | 8.06155228 | 6.6558E-30 |
| 5 | *cyp2k6* | 8.04849949 | 8.875E-15 |
| 6 | *soat2* | 7.97113062 | 7.111E-20 |
| 7 | *wu:fd46c06* | 7.5996353 | 2.3382E-56 |
| 8 | *si:ch211-220m17.4* | 7.4505096 | 1.7333E-12 |
| 9 | *CR855311.6* | 6.70373531 | 1.1751E-10 |
| 10 | *BX942813.1* | 6.57628174 | 3.1531E-09 |
| 11 | *chrne* | 6.49408558 | 2.0449E-13 |
| 12 | *PRSS35* | 6.34029035 | 1.5049E-08 |
| 13 | *BX248318.1* | 6.26372634 | 1.8779E-13 |
| 14 | *CR855311.4* | 6.24787071 | 7.4865E-09 |
| 15 | *amfrb* | 6.15766812 | 2.3779E-08 |
| 16 | *mtnr1al* | 5.93820758 | 2.0338E-07 |
| 17 | *aqp7* | 5.68958032 | 5.1375E-23 |
| 18 | *lrit3a* | 5.67875642 | 9.6784E-10 |
| 19 | *hsd11b2* | 5.53454033 | 1.1931E-13 |
| 20 | *aqp9b* | 5.43251912 | 2.7348E-06 |
| 21 | *lmod1a* | 5.43157199 | 1.6716E-06 |
| 22 | *tns1a* | 5.31570375 | 5.2367E-14 |
| 23 | *ugt5a2* | 5.25060333 | 1.3963E-18 |
| 24 | *phkg1b* | 5.09583142 | 4.1493E-20 |
| 25 | *CABZ01028768.1* | 5.08631544 | 2.1788E-09 |
| 26 | *hspb6* | 5.02313093 | 2.7874E-05 |
| 27 | *si:ch211-191a16.2* | 5.00961458 | 5.2017E-07 |
| 28 | *dio3b* | 4.96481276 | 8.2623E-06 |
| 29 | *dipk2ab* | 4.94643404 | 3.8081E-08 |
| 30 | *st3gal1l* | 4.91374375 | 2.3258E-05 |
| 31 | *me1* | 4.87259469 | 3.5874E-09 |
| 32 | *tg* | 4.64910663 | 5.3144E-14 |
| 33 | *zgc:92137* | 4.60175734 | 1.7141E-25 |
| 34 | *si:dkey-285b23.4* | 4.57523192 | 1.0474E-07 |
| 35 | *epha3* | 4.56245331 | 1.6346E-09 |
| 36 | *BX005064.1* | 4.55710583 | 1.1382E-05 |
| 37 | *CABZ01074397.1* | 4.49453907 | 7.1269E-15 |

**S3 Table Continued**

| Number based on Log2 fold change | Gene name | Log2 fold change | Adjusted p value |
|---|---|---|---|
| 38 | *si:ch211-150d5.3* | 4.46839097 | 1.4784E-06 |
| 39 | *slc12a10.3* | 4.46759273 | 0.0001258 |
| 40 | *fat2* | 4.43588616 | 6.6753E-05 |
| 41 | *il7r* | 4.37813055 | 6.3616E-06 |
| 42 | *sostdc1a* | 4.33825215 | 2.3073E-05 |
| 43 | *sox17* | 4.32621075 | 0.0005706 |
| 44 | *rab3c* | 4.25775534 | 0.00077508 |
| 45 | *si:dkey-71h2.2* | 4.16264725 | 1.6347E-11 |
| 46 | *ctrb1* | 4.16218319 | 1.5811E-20 |
| 47 | *crygm3* | 4.15624442 | 0.00105955 |
| 48 | *ela2* | 4.09857915 | 1.9526E-18 |
| 49 | *dpp6b* | 4.09364009 | 1.311E-08 |
| 50 | *gdf7* | 4.08951513 | 0.00160132 |
| 51 | *ddc* | 4.07206341 | 5.2061E-07 |
| 52 | *tmigd1* | 4.05886943 | 0.00160132 |
| 53 | *ins* | 4.0442376 | 0.00058318 |
| 54 | *spag6* | 4.0318073 | 0.00076682 |
| 55 | *cel.2* | 4.03180631 | 1.001E-16 |
| 56 | *DHRS11* | 4.01704462 | 3.8562E-14 |
| 57 | *nkx2.3* | 4.00022487 | 0.00175811 |
| 58 | *trim35-10* | 3.99780178 | 5.218E-05 |
| 59 | *ihha* | 3.99463338 | 2.5125E-05 |
| 60 | *col14a1b* | 3.97885055 | 0.0001919 |
| 61 | *c7b* | 3.93724042 | 4.8162E-09 |
| 62 | *si:dkey-266f7.9* | 3.93250088 | 0.00159986 |
| 63 | *pcxb* | 3.91745372 | 1.1931E-13 |
| 64 | *slc38a3a* | 3.87739216 | 2.3694E-07 |
| 65 | *zmp:0000000650* | 3.87705327 | 0.00296071 |
| 66 | *zgc:136461* | 3.87230106 | 1.2075E-07 |
| 67 | *amy2a* | 3.87218881 | 6.9627E-21 |
| 68 | *si:ch211-246e12.3* | 3.86744969 | 0.00267014 |
| 69 | *ctrl* | 3.85358994 | 9.2589E-18 |
| 70 | *apof* | 3.85328475 | 8.5537E-14 |
| 71 | *ela2l* | 3.8389998 | 8.0074E-14 |
| 72 | *CU469568.2* | 3.8380969 | 4.5061E-07 |
| 73 | *ugt5a4* | 3.83739905 | 7.5169E-08 |
| 74 | *cyp3a65* | 3.83221644 | 2.2291E-07 |
| 75 | *cela1.5* | 3.82188785 | 4.8426E-12 |
| 76 | *cpa4* | 3.80213184 | 2.3069E-06 |
| 77 | *gcga* | 3.79973222 | 1.2315E-05 |

**S3 Table Continued**

| Number based on Log2 fold change | Gene name | Log2 fold change | Adjusted p value |
|---|---|---|---|
| 78 | *stk24a* | 3.79940557 | 8.3244E-05 |
| 79 | *prss1* | 3.79774569 | 2.3667E-18 |
| 80 | *si:ch211-266k22.6* | 3.78158496 | 0.00573631 |
| 81 | *cel.1* | 3.76552094 | 2.5288E-14 |
| 82 | *abcg2c* | 3.74036036 | 3.4991E-14 |
| 83 | *ugt5a1* | 3.73156129 | 0.00184259 |
| 84 | *c6ast3* | 3.72415875 | 4.0719E-15 |
| 85 | *mettl25* | 3.71768857 | 4.1001E-28 |
| 86 | *si:dkey-85k7.10* | 3.67149845 | 0.00508226 |
| 87 | *si:ch211-255i20.3* | 3.66923139 | 0.00561862 |
| 88 | *gch1* | 3.65963182 | 2.969E-06 |
| 89 | *cela1.1* | 3.6582973 | 1.1996E-18 |
| 90 | *si:ch211-240l19.6* | 3.64974117 | 9.0669E-10 |
| 91 | *fosl1a* | 3.64457454 | 9.2537E-05 |
| 92 | *hsp70l* | 3.63849194 | 0.0004497 |
| 93 | *si:dkey-285b23.3* | 3.63786224 | 5.0828E-07 |
| 94 | *cela1.6* | 3.63738924 | 1.2329E-16 |
| 95 | *myo3a* | 3.63324954 | 0.00017664 |
| 96 | *mgp* | 3.63269734 | 0.00468451 |
| 97 | *gch2* | 3.62937132 | 2.8062E-07 |
| 98 | *dnase1* | 3.62908863 | 2.6815E-05 |
| 99 | *BX323793.1* | 3.62830803 | 0.00421167 |
| 100 | *znrf1* | 3.62660647 | 1.4693E-06 |
| 385 | *thbs1a* | 2.2095974 | 2.29392E-05 |
| 461 | *ezra* | 2.0262628 | 8.83399E-05 |

**S4 Table. Genes upregulated in *podxl*$^{Ex1(p)\_Ex7\Delta}$ mutants are enriched in zebrafish HSCs.**

We examined genes that were significantly upregulated in *podxl*$^{Ex1(p)\_Ex7\Delta}$ mutants (log2fold change> 2, padj<0.05) alongside three publicly available datasets for HSC-enriched genes. Genes meeting criteria for HSC enrichment and/or upregulation, marked with "x", in at least two of these four datasets are included in this table.

| **Gene** | ***podxl* mutant** | **Yin** (Yin et al. 2012) | **Morrison** (Morrison et al. 2022) | **Spanjaard** (Spanjaard et al. 2018) |
|---|---|---|---|---|
| *ablim1a* | x | x | | |
| *acta2* | x | x | | |
| *actb2* | x | | | x |
| *anxa1a* | | x | | x |
| *anxa2a* | | x | | x |
| *atp1a3b* | x | x | | |
| *cavin2b* | x | | | x |
| *col1a1b* | x | x | | |
| *cpa2* | x | x | | |
| *fabp11a* | | x | x | |
| *fosl2* | x | | x | |
| *frzb* | x | | | x |
| *gcga* | x | x | | |
| *hsp70l* | x | | x | |
| *hspb1* | | x | x | |
| *ifitm1* | | | x | x |
| *jun* | x | | x | |
| *kdrl* | | x | x | |
| *krt4* | x | x | | x |
| *krt8* | | | x | x |
| *krt94* | | | x | x |
| *mb* | | x | x | |
| *mdka* | | x | | x |
| *mmp2* | | x | | x |
| *mvp* | x | | x | |
| *pmp22b* | | x | x | |
| *podxl* | | x | | x |
| *rnd3b* | x | x | | |
| *s100a10b* | | x | | x |
| *slc43a3b* | x | | | x |
| *sostdc1a* | x | x | | |
| *tagln* | x | x | | |

| | | | | |
|---|---|---|---|---|
| *tmem88b* | | x | | x |
| *tmsb1* | x | | | x |
| *vldlr* | x | x | | |
| *zgc:154093* | x | x | | |

**S5 Table. Expression of genes highly similar to *podxl* in $podxl^{Ex1(p)\_Ex7\Delta}$ mutants.**

| Gene name | Query cover (percentage) | E value | Percent identity | Log 2 fold change | Adjusted p-value |
|---|---|---|---|---|---|
| LOC108179520 | 10% | 4E-83 | 80.73% | Not detected | Not detected |
| Magixa | 5% | 3E-59 | 86.18% | -0.53697307 | 0.71511715 |
| Endo | No significant similarity | NA | NA | Not detected | Not detected |

**S6 Table. Expression of genes somewhat similar to *podxl* in $podxl^{Ex1(p)\_Ex7\Delta}$ mutants.**

| Gene name | Query cover (percentage) | E value | Percent identity | Log 2 fold change | Adjusted p-value |
|---|---|---|---|---|---|
| LOC108179520 | 14% | 1E-126 | 75.49% | Not detected | Not detected |
| Psd2 | 14% | 2E-98 | 72.83 | 0.56784281 | 0.17828786 |
| Synm | 11% | 8E-67 | 79.22 | 0.50306331 | 0.80274464 |
| Xpa | 5% | 1E-58 | 83.26% | -0.57652444 | 0.71066531 |
| Magixa | 5% | 3E-65 | 86.18% | -0.53697307 | 0.71511715 |
| Washc3 | 5% | 3E-52 | 81.65% | -0.64157183 | 0.33876005 |
| Zfyve9a | 5% | 9E-60 | 84.33% | -0.63260206 | 0.29769286 |
| Pdzd2 | 5% | 9E-60 | 87.11% | -0.84252339 | 0.34490123 |
| Clpxa | 5% | 2E-56 | 83.18% | 0.72980566 | 0.2934432 |
| Usp16 | 5% | 5E-56 | 85.64% | -0.57259874 | 0.70027044 |
| Ywhaqb | 5% | 7E-55 | 83.10% | 0.38869663 | 0.62509361 |
| Endo | No significant similarity | NA | NA | Not detected | Not detected |

**S7 Table: Expression of genes surrounding *podxl* on chromosome 4 in *podxl*$^{Ex1(p)\_Ex7\Delta}$ mutant adult livers versus wildtype control sibling livers.** We examined all coding genes within 700 kbp of the *podxl* start site.

| Gene ID<br>Transcript ID | Gene Symbol | log2 Fold Change | p-value (adj) | Start site | Kbp From Start Site to Podxl Start site |
|---|---|---|---|---|---|
| ENSDARG00000019396<br>ENSDART00000017180.8 | *rergla* | 1.8516 | 0.0858 | 12388312 | -637 |
| ENSDARG00000058244<br>ENSDART00000081089.4 | *il17ra1a* | -0.2512 | 0.6772 | 12322993 | -572 |
| ENSDARG00000041665<br>ENSDART00000061070.8 | *mkrn1* | 0.2700 | 0.5947 | 12292298 | -541 |
| ENSDARG00000007639<br>ENSDART00000022646.7 | *cnot4b* | 0.2379 | 0.6119 | 12277917 | -527 |
| ENSDARG00000020015<br>ENSDART00000048675.6 | *mrps33* | -0.1617 | 0.6958 | 12104192 | -353 |
| ENSDARG00000017661<br>ENSDART00000048391.10 | *Braf* | -0.5752 | 0.1793 | 12101841 | -351 |
| **ENSDARG00000045768<br>ENSDART00000130692.4** | ***cry1a*** | **0.9196** | **0.0007** | **12013736** | **-263** |
| ENSDARG00000063255<br>ENSDART00000092250.7 | *btbd11a* | 0.5731 | 0.4640 | 12006752 | -256 |
| ENSDARG00000005891<br>ENSDART00000014153.6 | *cyb5r3* | -0.1380 | 0.7412 | 11810647 | -60 |
| **ENSDARG00000031228<br>ENSDART00000102301.5** | ***podxl*** | **-5.2161** | **<0.0001** | **11751037** | **0** |
| ENSDARG00000025576<br>ENSDART00000137736.4 | *mkln1* | 0.2621 | 0.3673 | 11696005 | 55 |
| **ENSDARG00000032765<br>ENSDART00000049066.6** | ***net1*** | **1.4129** | **0.0006** | **11580677** | **170** |
| ENSDARG00000045765<br>ENSDART00000140954.2 | *asb13a.2* | 0.9235 | 0.2386 | 11483487 | 268 |
| ENSDARG00000079698<br>ENSDART00000019458.9 | *asb13a.1* | 0.7465 | 0.1985 | 11479823 | 271 |
| ENSDARG00000005451<br>ENSDART00000008584.5 | *gdi2* | 0.0851 | 0.7210 | 11464358 | 287 |

| | | | | | |
|---|---|---|---|---|---|
| ENSDARG00000003822 ENSDART00000037600.7 | *ankrd16* | 0.6555 | 0.2828 | 11458235 | 293 |
| **ENSDARG00000019235 ENSDART00000051792.6** | ***sema3aa*** | **2.5507** | **0.0000** | **11311708** | **439** |
| ENSDARG00000001781 ENSDART00000138661.3 | *tspan11* | 0.0457 | 0.9494 | 11175566 | 575 |
| ENSDARG00000045758 ENSDART00000142389.4 | *cracr2ab* | -0.1421 | 0.8937 | 11135048 | 616 |
| ENSDARG00000045755 ENSDART00000140362.3 | *ccdc59* | -0.7998 | 0.0665 | 11053778 | 697 |
| **ENSDARG00000045754 ENSDART00000067262.6** | ***mettl25*** | **3.7177** | **0.0000** | **11053278** | **698** |

**S8 Table. gRNAs for generation of *podxl* mutants**

| PCR primers for genotyping and fragment analysis | sgRNA 1 | sgRNA 2 |
|---|---|---|
| *podxl*$^{Ex1,-5bp\Delta}$ mutant | TACGATGATTGTCCACGTGATGG | |
| *podxl*$^{Ex1(p)\_Ex7\Delta}$ mutant | TACGATGATTGTCCACGTGATGG | GTTAGGAGGTGAGTGTTACCTGG |
| *podxl*$^{Ex5\_Ex7\Delta}$ mutant | CAGATCTGTAGCTCATGCGGGGG | GTTAGGAGGTGAGTGTTACCTGG |
| *podxl*$^{-319\_Ex1(p)\Delta}$ mutant | CAGATCTGTAGCTCATGCGGGGG | TACGATGATTGTCCACGTGATGG |
| *podxl* $^{-194\_Ex7\Delta}$ mutant | AAATCCCCGGACCCTCAGGGTGG | GTTAGGAGGTGAGTGTTACCTGG |

**S9 Table. PCR primers for genotyping *podxl* mutants**

| Podxl mutant, allele target | Forward primer | Reverse primer | Band size | Annealing temp |
|---|---|---|---|---|
| *podxl*$^{Ex1,-5bp\Delta}$ mutant, Mutant allele | GACTGAACGCGGAGAATCTG | CGTCGTATATGAAGGTAAACTCACC | 73 bp | 68.3 |
| *podxl*$^{Ex1,-5bp\Delta}$ mutant, WT allele | GACTGAACGCGGAGAATCTG | CGTCGTATATGAAGGTAAACTCACC | 78 bp | 68.3 |
| *podxl*$^{Ex1(p)\_Ex7\Delta}$ mutant, Mutant allele | CAAGACGAAAAGCGGAACCG | TGCCATTCTTGGAGTTCCTTCA | 155 bp | 58 |
| *podxl*$^{Ex1(p)\_Ex7\Delta}$ mutant, WT allele | TGCCATTCTTGGAGTTCCTTCA | TGCCATTCTTGGAGTTCCTTCA | 450 bp | 58 |
| *podxl*$^{Ex5\_Ex7\Delta}$ mutant, Mutant allele | TTTCATCCTAAAATTCACCTGCACA | GTGTTCCTTTCCGGGTCCAT | 394 bp | 58 |
| *podxl*$^{Ex5\_Ex7\Delta}$ mutant, WT alllele | TACAGAACAATGGAGCCAGAGG | AATACGAGGAAAAGTGATTCCTCCA | 300 bp | 58 |
| *podxl*$^{-319\_Ex1(p)\Delta}$ mutant, Mutant allele | TGGGACCAGCCCTCTGTG | TAAACCCAGCACATGTCCCTC | 163 bp | 58 |
| *podxl*$^{-319\_Ex1(p)\Delta}$ mutant, WT allele | CACACCAAAACTTTGCCCCC | TAAACCCAGCACATGTCCCTC | 373 bp | 58 |
| *podxl*$^{-194\_Ex7\Delta}$ mutant, Mutant allele | AGAGCCAGTCACTCCAGACT | AAAGTGTGTTCCTTTCCGGGT | 423 bp | 58 |
| *podxl*$^{-194\_Ex7\Delta}$ mutant, WT allele | TGGGACCAGCCCTCTGTG | TAAACCCAGCACATGTCCCTC | 492 bp | 58 |
